# The differential effect of optogenetic serotonergic manipulation on sustained motor actions and stationary waiting for future rewards in mice

**DOI:** 10.1101/2024.05.17.594118

**Authors:** Masakazu Taira, Kayoko W. Miyazaki, Katsuhiko Miyazaki, Jianning Chen, Shiho Okitsu-Sakurayama, Anupama Chaudhary, Mika Nishio, Tsukasa Miyake, Akihiro Yamanaka, Kenji F. Tanaka, Kenji Doya

**Author notes:** **Correspondence:** Masakazu Taira, Kenji Doya.

## Abstract

Serotonin is an essential neuromodulator that affects behavioral and cognitive functions. Previous studies have shown that activation of serotonergic neurons in the dorsal raphe nucleus (DRN) promotes patience to wait for future rewards. However, it is still unclear whether serotonergic neurons also regulate persistence to act for future rewards. Here we used optogenetic activation and inhibition of DRN serotonergic neurons to examine their effects on sustained motor actions for future rewards. We trained mice to perform stationary waiting and repeated lever-pressing tasks with variable reward delays and tested effects of optogenetic activation and inhibition of DRN serotonergic neurons on task performance. Interestingly, in the lever-pressing task, mice tolerated longer delays as they repeatedly pressed a lever than in the stationary waiting task, suggesting that lever-pressing actions may not simply be costly, but may also be subjectively rewarding. Optogenetic activation of DRN serotonergic neurons prolonged waiting in the stationary waiting task, consistent with previous studies, but it did not affect lever pressing time or numbers. While optogenetic inhibition decreased waiting, it did not affect lever pressing time or numbers. In generalized linear model analysis that incorporated the time during each session and the number of sessions, however, optogenetic activation negatively affected the number and the speed of lever pressing. These results revealed that the necessity of motor actions may increase motivation for delayed rewards and that DRN serotonergic neurons more significantly promote stationary waiting rather than persistent motor actions for future rewards.

## 1 Introduction

Serotonin (5-HT) is an important neuromodulator involved in multiple biological functions, including emotion (Cools et al., 2008), motivation (Dayan and Huys, 2009), motor activity (Ranade and Mainen, 2009), and decision making (Homberg, 2012). 5-HT projections originate from raphe nuclei located in the midbrain. Among 9 raphe nuclei, the dorsal raphe nucleus (DRN) densely projects to the forebrain and DRN 5-HT neurons regulate reward-based learning and decision making (Miyazaki et al., 2012a, Doya et al., 2021).

Previous computational studies based on the reinforcement learning framework, proposed that 5-HT controls the temporal discount factor and that activation of 5-HT neurons increases the relative importance of future rewards over immediate rewards (Doya, 2002, Schweighofer et al., 2007). In support of this hypothesis, a series of experimental studies has been done using delayed reward tasks, showing increased 5-HT transmission while rats were waiting for delayed rewards (Miyazaki et al., 2011a, Miyazaki et al., 2011b). Furthermore, pharmacological inhibition of DRN 5-HT neurons increased premature abandonment of delayed rewards (Miyazaki et al., 2012b) and optogenetic activation of those neurons prolonged the time spent for waiting for delayed rewards, establishing a causal relationship between DRN 5-HT neurons and patience in waiting for future rewards (Miyazaki et al., 2014, Fonseca et al., 2015, Miyazaki et al., 2018, Miyazaki et al., 2020). While these studies examined the role of DRN 5-HT neurons in passive waiting to obtain future rewards, how these neurons regulate active behavior to obtain future rewards has not been well studied, except in the context of patch-leaving decision-making (Lottem et al., 2018).

To examine the role of DRN 5-HT neurons in sustained motor actions, we trained mice to perform an operant conditioning task that requires variable numbers of lever-presses, and we tested the effect of optogenetic activation and inhibition of DRN 5-HT neurons on motor actions. For comparison, we also tested the effect of optogenetic manipulation of DRN 5-HT neurons in a stationary waiting task. We found that optogenetic activation of DRN 5-HT neurons prolonged waiting, but that it did not affect the duration or the number of lever presses for future rewards. While optogenetic inhibition reduced the waiting time, it had no effect on the duration or the number of lever presses for future rewards. Further analyses using generalized linear models (GLMs) revealed a negative effect of DRN 5-HT activation on the number and the speed of lever pressing. These results suggest that DRN 5-HT neurons regulate two types of behaviors for future rewards in different ways.

## 2 Materials and Methods

### 2.1 Animals

All experimental procedures were performed in accordance with guidelines established by the Okinawa Institute of Science and Technology Experimental Animal Committee. For the optogenetic activation experiment, we used Tph2-ChR2(C128S)-EYFP bi-transgenic mice. ChR2(C128S) is a step-type function opsin that remains activated by short pulses of blue light and deactivated by yellow light (Miyazaki et al., 2014). Eight Tph2-ChR2 mice were trained for the lever-pressing task before implantation of optic probes. Among them, four mice were first tested in the lever-pressing task and then the stationary waiting task, while the other four mice were tested in the opposite order. For controls, five Tph2-tTA transgenic mice were used. All Tph2-tTA mice were first tested in the lever-pressing task followed by the waiting task.

For optogenetic inhibition experiments, we used Tph2-ArchT-EGFP bi-transgenic mice. ArchT activates an inhibitory current in response to yellow light (Tsutsui-Kimura et al., 2017, Yoshida et al., 2019). We trained separate cohorts of Tph2-ArchT mice for different behavioral tasks. Five Tph2-ArchT mice were used for the stationary waiting task and the other six mice were used for the lever-pressing task. Tph2-tTA transgenic mice were used as a control group. Four Tph2-tTA mice were used for the stationary waiting task and five Tph2-tTA mice were used for the lever-pressing task. We generated bi-transgenic mouse lines as shown in Tanaka et al. (2012). All mice have the background of C57BL/6 mice. All mice were male and training started > 4 months of age.

All mice were housed individually at 24 ℃ on a 12:12 h light: dark cycle (lights on 07:00-19:00 h). All behavioral training and test sessions were performed during the light cycle, 5 days per week. Mice were deprived of access to food one day prior to the first training session and received daily food rations only during training and test sessions (approximately 2-3 g per day) during experimental days. Food was freely available during days off more than 15 hours before the next session. Mice could freely access water in their home cages.

### 2.2 Behavioral apparatus

All training and testing sessions were performed in operant boxes (Med-associates, 21.6 cm width x 17.8 cm depth x 12.7 cm height). Two 2.5-cm, square holes were located in the walls on opposite sides of the box. One hole was designated as the reward site, and was connected to a food dispenser delivering 20-mg food pellets, while the other hole was defined as a tone site. A retractable lever was positioned to the left of the reward site. One 2.8-W house light and one speaker were located above and to the upper right of the tone site, respectively. Hardware attached to operant boxes was controlled via MED-PC IV software (Med-associates).

### 2.3 Tone-food waiting task

#### 2.3.1 Task structure

We used the same behavioral task as reported in previous studies (Figure 1A(i); (Miyazaki et al., 2014, Miyazaki et al., 2018). In this task, mice could initiate a trial with a 0.3-s nose poke to the tone site, which triggered a 0.5-s tone. After hearing the tone, mice were required to continue poking their noses into the reward site. The required duration of the nose-poke was randomly chosen as 2, 6, 10 s, and infinity (reward omission) during each trial. Once mice could wait for the required time, a food pellet was delivered to the reward site. For the photoactivation experiment, mice could initiate the next trial just after a reward delivery or after leaving the reward site. Because the suppression efficacy of ArchT decreases without sufficient duration of intervals between light stimulation (Mattis et al., 2011), for the photoinhibition experiment, the house light was turned off for 30 s after the end of a trial and the next trial could be initiated once the house light was turned on again. One session consisted of 43 trials (5 trials x 2 photostimulation conditions x 4 delays + 3 trials with different photostimulation and delay conditions). Three sessions were performed on a testing day. Before the testing sessions, mice were trained for 2 h, five days per week, and it took 2 weeks or less for mice to learn the task.

**Figure 1.**
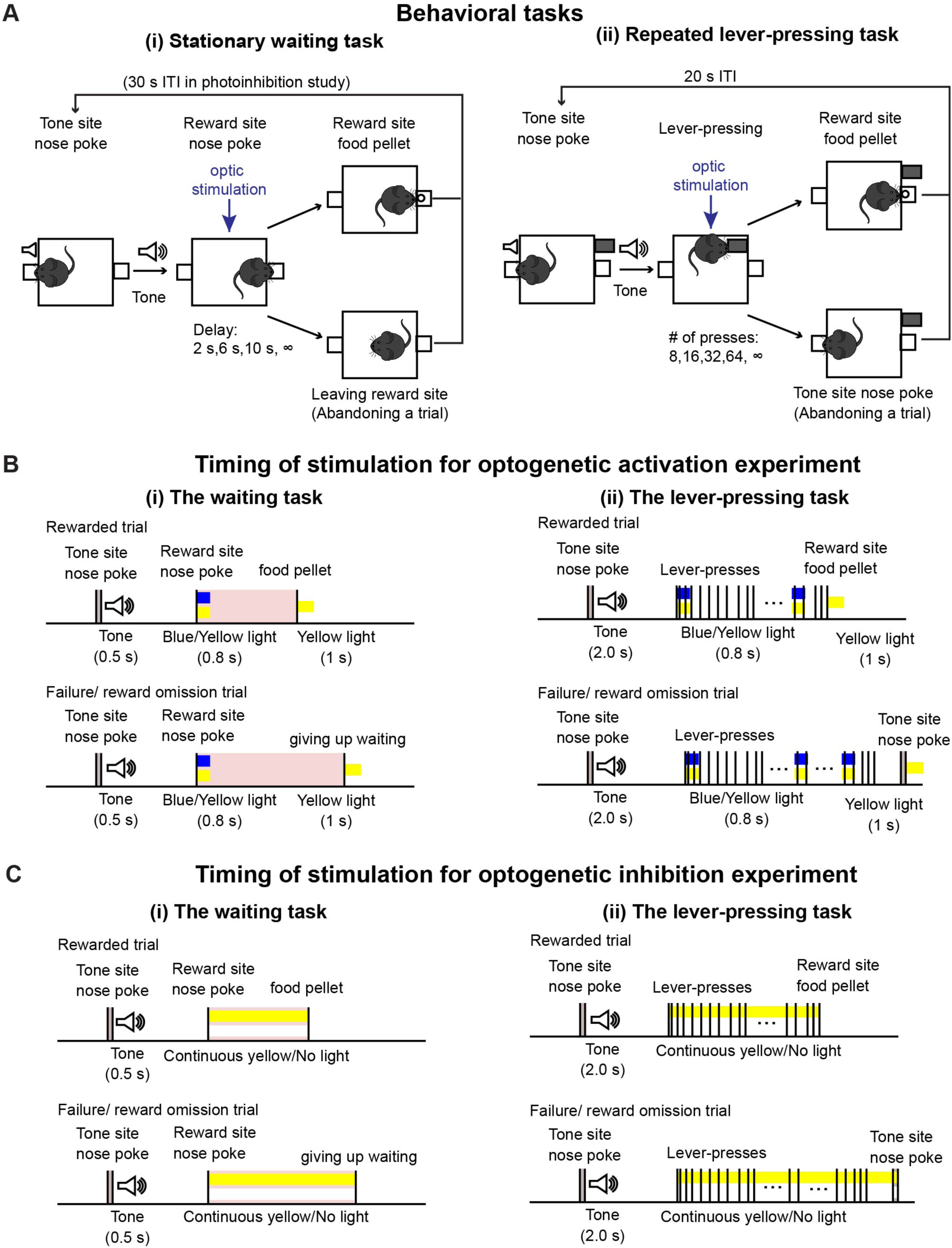
Behavioral tasks. (A) Schematic drawing of (i) the stationary waiting task and (ii) the repeated lever-pressing task. (B) Time sequence of rewarded and failure/reward omission trials with optic stimulation for optogenetic activation experiments during (i) the waiting task and (ii) the lever-pressing task. (C) Time sequence of rewarded and failure/reward omission trials with optic stimulation for optogenetic inhibition experiments during (i) the waiting task and (ii) the lever-pressing task.

#### 2.3.2 Timing of optic manipulation

During testing sessions, 470-nm blue or 590-nm yellow light stimulation was given, generated by an LED light source (Doric Lenses). Timing of stimulation was effected by TTL pulses controlled with MED-PC IV software. For the optogenetic activation experiment (Figure 1B(i)), in half the trials (selected at random), blue light stimulation was applied for activation, and in the other half, yellow light stimulation was given as a control. A 0.8-s blue/yellow light pulse was given when mice first poked their noses into the reward site. At the end of the trial, either when the mice waited until the end for the required delay or when they left the reward site, a 1-s yellow light pulse was given to reset photoactivation. The intensity of blue and yellow light at the tip of the optical fibers was 1.6-2.0 mW and 1.1-2.0 mW, respectively. For the optogenetic inhibition experiment (Figure 1C(i)), in half the trials, yellow light stimulation was applied for inhibition, while no light was applied for control trials in the remaining trials. In the inhibition trials, continuous yellow light was applied from the onset of waiting until the end of a trial. Intensities of yellow light at the tip of the optical fibers were 2.8-3.2 mW.

### 2.4 Variable number lever-pressing task

#### 2.4.1 Task structure

Task structure is described in Figure 1A(ii). After the house light was turned on, mice could initiate a trial by poking their noses into the tone site for 0.3 s. The 0.3-s nose poke triggered a speaker to generate a 2-s tone, after which a retractable lever was presented. The number of lever-presses required was randomly chosen as 8, 16, 32, 64, and infinity (reward omission) during each trial. Once mice pressed the lever the required number of times, the lever was withdrawn, and 1 s after lever withdrawal, a food pellet was delivered to the reward site. Alternatively, mice could abandon the trial with a 0.3-s nose poke into the tone site. After reward delivery or abandonment of the trial, a 15-s inter-trial interval was inserted, which was indicated by turning off the house light. After the 15-s inter-trial interval, mice could initiate the next trial. One testing session consisted of 53 trials (5 trials x 2 photostimulation conditions x 5 press number conditions + 3 trials with different photostimulation and press conditions). Two sessions were performed on a given testing day.

#### 2.4.2 Timing of optic manipulation

For the optogenetic activation experiment (Figure 1B(ii)), in half the trials (selected at random), blue light stimulation was applied for activation, and in the other half, yellow light stimulation was given as a control. A 0.8-s blue/yellow light pulse was given when the mice started to press the lever and repeated at 20-s intervals. At the end of a trial, either when the mice pressed the lever the required number of times, or when they abandoned the trial, a 1-s yellow light pulse was given. For the optogenetic inhibition experiment (Figure 1C(ii)), in half the trials, yellow light stimulation was applied for inhibition, while no light was applied for control trials in the remaining trials. In the inhibition trials, continuous yellow light was applied from the onset of lever-pressing until the end of a trial. We used the same LED light source and light intensity from the testing sessions of the waiting task described in a previous section.

#### 2.4.3 Training procedures

Before testing sessions commenced, mice were trained to perform the lever-pressing task using the following schedule. Training took approximately 3 weeks.

All training sessions were performed either until mice earned 100 food pellets or by 2 h, whichever came first. In order to habituate mice to the behavioral apparatus, they were first trained to poke their noses into the reward site to obtain a food pellet. Then they were trained to press a lever once to acquire a food pellet. Once mice could get more than 80 food pellets, they were trained to press a lever 3 times to obtain a food pellet. While the number of lever-presses required was progressively increased from 3, 5, 7, 10, 16, and to 32 times, mice were trained to learn that the lever was presented after a tone was generated. After the association between tone and lever presentation was established, mice were trained to poke their noses to generate a tone for lever presentation. Training was complete when mice could get more than 80 rewards in a training session, during which they were required to press the lever 32 times after initiating a trial using nose pokes.

### 2.5 Surgical procedure for optic probe implantation

After training for the lever-pressing task was completed, a craniotomy was performed to implant an optic probe (400 μm diameter, 0.48 NA, 4 mm length, Doric) above the DRN. Mice were anesthetized with isoflurane (3% for induction and 1-1.5% during surgery). Mice were placed on a stereotaxic stage and their heads were fixed with ear bars. Then the skull was exposed with a blade, and a hole was made with a drill. Once the brain was exposed through the hole, the dura was removed using the tip of a needle, and the optic probe was lowered above the DRN through the hole (from the bregma: posterior, −4.6mm; lateral, 0 mm; ventral, −2.6 mm). Light-sensitive adhesive and dental cement was applied to the skull to fix the implanted optic probe. Mice were placed back in their home cages for recovery. At least one week after the surgery, we started to retrain the mice for the behavioral task, and then commenced testing sessions.

### 2.6 Histological confirmation of the implantation site

After the behavioral tests, mice were deeply anesthetized with 100 mg/kg sodium pentobarbital i.p. and perfused with saline or PBS followed by 4% PFA/PB or 4% PFA/PBS. Brains were removed immediately after perfusion and immersed in fixative solution overnight. Then, 50-μm coronal slices were sectioned using a vibratome (VT1000S, Leica) and the implantation site of optic probes was confirmed according to the mouse brain atlas (Franklin, 2008).

### 2.7 Immunohistochemistry

Brain slices were incubated with primary antibodies for 1-2 nights. Slices were then rinsed with PBS and incubated with secondary antibodies for 1-2 nights. After incubation and rinsing, slices were mounted on slide glasses. Fluorescent images (Figure 2B) were acquired using a spinning disc confocal microscope (Nikon). As primary antibodies, we used anti-Tph (1:250, sheep polyclonal, Merck Millipore, AB1541) and anti-GFP (1:500, chicken polyclonal, Abcam, ab13970) as markers for 5-HT neurons and ChR2-EYFP or ArchT-EGFP neurons, respectively. For secondary antibodies, anti-sheep and anti-chicken antibodies conjugated with Alexa flour 594, and 488, respectively, were used. Antibodies were diluted in staining buffer containing 10mM HEPES, 20mM NaCl, and 10% Triton-X100. The pH of the staining buffer was adjusted to 7.4 in advance.

**Figure 2.**
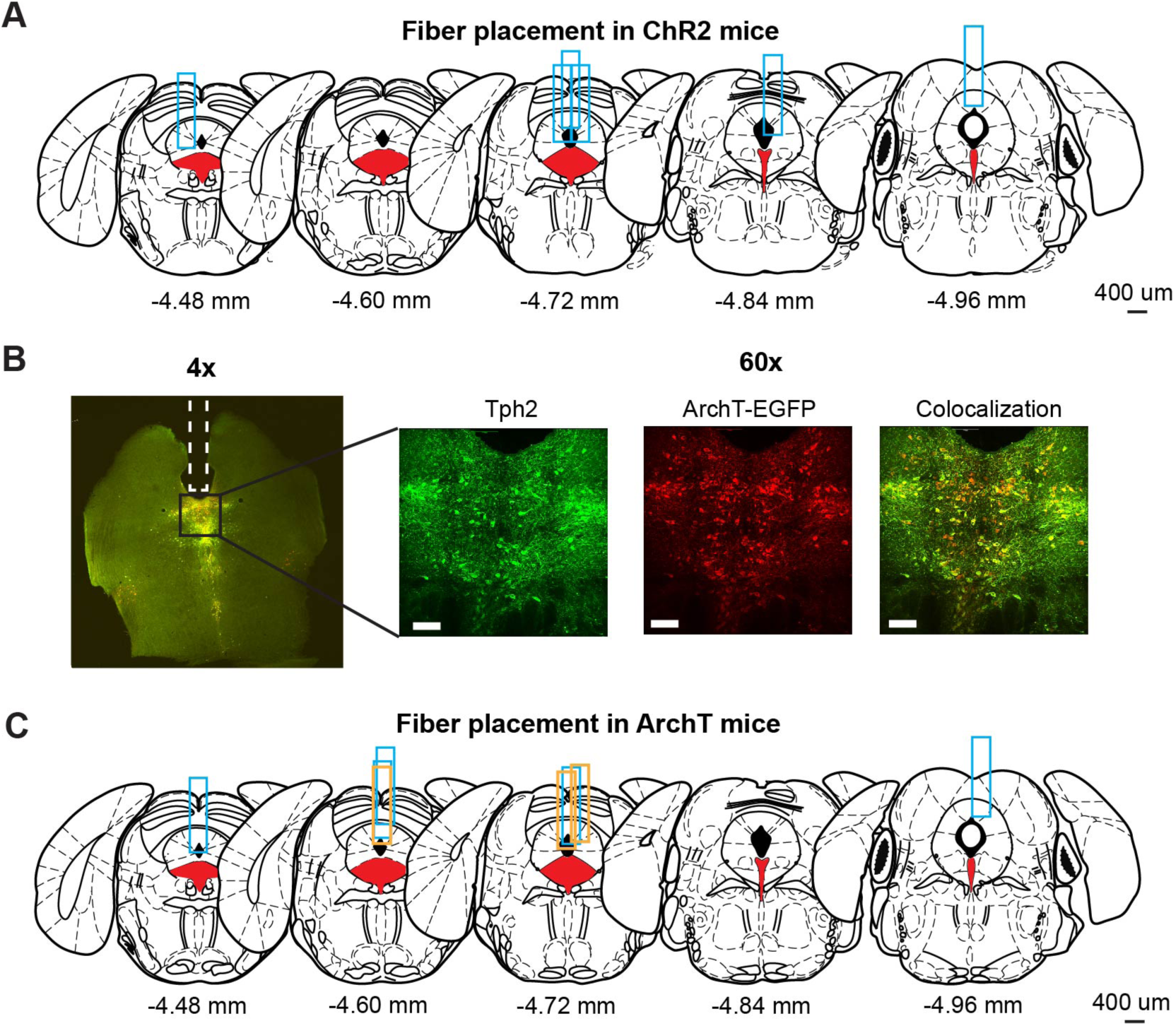
Optogenetic manipulation of DRN 5-HT neural activities. (A) The implantation site of optic probes for ChR2 mice. Coronal views of the mouse brain are adapted from Franklin (2008). Red filled areas indicate the DRN. Blue rectangles indicate tracks of implanted optic fibers. (B) Fluorescence images from ArchT mice. The left image (4x) shows fluorescent signal from Tph2 (red) and ArchT-EGFP (green) in a coronal brain section including DRN. Magnified images (60x) indicate the florescent signal of Tph2 (Left), ArchT-EGFP (Middle), and co-localization of the two signals (Right). Scale bars indicate 100 μm. (C) The implantation site of optic probes for ArchT mice. Coronal views of the mouse brain are adapted from Franklin (2008). Red filled areas indicate the DRN. Blue and orange rectangles indicate the track of optic fibers in the ArchT mice used for lever-pressing and waiting tasks, respectively.

### 2.8 Behavioral parameters and statistical analysis

In the lever-pressing task, the successful trial rate for 8-, 16-, 32-, and 64-press trials was calculated by dividing the number of rewarded trials by the total number of trials. In omission trials, the number of lever-presses, the time spent lever-pressing, which was defined as the time elapsed from the first lever-press to the last, and the time to abandon an omission trial, which was defined as the duration between the last lever-press and a nose poke to terminate the trial, were measured. To examine action vigor, inter-press intervals (IPIs), and intervals between successive lever-presses, were measured. IPIs longer than 5 s were defined as long IPIs and below 5 s as short IPIs. In the waiting task, the duration of maintaining a nose poke was measured in omission trials. Behavioral parameters were calculated using custom-written programs in MATLAB.

Statistical tests for summary statistics were selected based on whether data exhibited normality and homogeneity of variance, assessed by the Shapiro-Wilk test and Levene test, respectively. If data satisfied these assumptions, we used paired t-tests for within-group comparison and unpaired t-tests for group comparisons. If not, we used the Wilcoxon signed-rank test for within-subject comparisons and the Mann Whitney U-test for group comparisons. Statistical analysis was performed using Python. For analysis of short IPIs, we applied repeated measures analysis of variance (ANOVA) using SPSS.

### 2.9 Generalized linear model (GLM) analysis

To examine the effect of optogenetic manipulation while taking into account multiple task-relevant variables and individual variability, we constructed GLMs and fit them to behavioral data. Response variables were the number of lever-presses and the short IPIs. For the number of lever-presses, we used logarithm link function and assumed a Poisson distribution. For short IPIs, we assumed a normal distribution and used an identity link function. Fixed effect terms were intercept, manipulation, elapsed time and session index, and the random effect term was the intercept for individuals. Manipulation was coded as 0 for control trials, i.e., yellow light/no-light trials in activation/inhibition studies, and 1 for intervention trials, i.e., blue light/yellow light trials in activation/inhibition studies. Elapsed time was how many seconds elapsed since the start of the session until a nose-poke to initiate an omission trial (up to 7200 s). Session index denoted how many sessions the subject had experienced across experimental periods (up to 8 sessions). GLM analysis was performed with the *fitglme* function in MATLAB. t-statistics and p-values were calculated to test the null hypothesis that the estimated regression coefficients were equal to zero.

## 3 Results

### 3.1 Histological confirmation of opsin expression and probe locations

We used eight Tph2-ChR2(C128S)-EYFP bi-transgenic mice (hereafter, referred to as ChR2 mice) to selectively activate DRN 5-HT neurons following a pulse of blue light, as in previous studies (Miyazaki et al., 2014, Miyazaki et al., 2018). Six of the eight ChR2 mice were sacrificed to confirm the implantation site of the optic probes. We could not perfuse the other two ChR2 mice because they died before the process. Although the site varied along the anterior-posterior axis, all probes examined were located above the DRN (Figure 2A).

To selectively inhibit DRN 5-HT neurons, we prepared eleven Tph2-ArchT-EGFP bi-transgenic mice (hereafter, referred to as ArchT mice). These transgenic mice selectively express ArchT, a light-sensitive proton pump, in 5-HT neurons. When sensing yellow light, ArchT induces efflux of H^+^ and inhibits neural activities. To confirm selective expression of ArchT, we performed a histological experiment to compare cells expressing EGFP and 5-HT neurons identified by Tph2 immunohistochemistry in three ArchT mice that were not used for the photoinhibition experiment. In nine slices from the three ArchT mice, 1058 Tph-positive cells were found. Of these cells, 72.5% were also ArchT-EGFP-positive. On the other hand, there were very few Tph-negative, but ArchT-EGFP-positive cells, suggesting that Tph2-ArchT bi-transgenic mice selectively expressed ArchT in 5-HT neurons. We quantitatively measured Tph and ArchT cells in five slices from the three ArchT mice that had experienced optogentic inhibition test sessions for 4 weeks of experiments. Percentages of Tph-positive and ArchT-positive cells were 69.8%, which was comparable to non-stimulated samples. Numbers of Tph-positive and ArchT-positive cells in stimulated samples were not significantly different from those in non-stimulated samples (stimulated vs non-stimulated: 57.60 ± 24.00 and 85.11 ± 11.19, t_12_ = 1.82, p = 0.0829, unpaired t-test). In separate cohorts of ArchT mice used for the present studies, we found abundant expression of ArchT. Colocalization of ArchT and Tph was confirmed in ArchT mice (Figure 2B), suggesting that optogenetic inhibition did not induce severe damage in 5-HT neurons after the experimental sessions. We also confirmed the implantation site of optic probes in eight of eleven ArchT mice used in lever-pressing and waiting tasks. We could not perfuse the other three mice for various reasons: one that died before perfusion, another that was used in another experiment and the third for unexpected removal of the probe before perfusion. All optic probes confirmed were implanted above the DRN in ArchT mice (Figure 2C). Data from those three mice were also included, since their data showed trends similar to those from the other eight mice.

### 3.2 Optogenetic activation of DRN 5-HT neurons

#### 3.2.1 Activation of DRN 5-HT neurons prolonged stationary waiting for future rewards

Mice were trained to perform two operant conditioning tasks: a stationary waiting task and a repeated lever-pressing task. In order to optogenetically activate DRN 5-HT neurons, a 0.8-s blue/yellow light pulse was given at the onset of a trial and a 1-s yellow light pulse was applied at the end thereof (Figure 1B(i) for the waiting task and 1B(ii) for the lever-pressing task). In the lever-pressing task, short blue/yellow pulses were applied every 20 s to refresh photostimulation (Figure 1B(ii)).

To confirm the effectiveness of optogenetic activation, we examined whether the optogenetic activation protocol used here affected waiting behaviors for delayed rewards of 2 s, 6 s, 10 s or infinity (reward omission). (Figure 3A). Optogenetic activation with blue light significantly increased waiting duration during omission trials in ChR2 mice (t_7_ = 6.31, p = 0.00040, paired t-test; Figure 3B). The change of duration in ChR2 was significantly larger than that in control mice (t_11_ = 4.98, p = 0.00042, unpaired t-test; **Figure** 3B-D). This result was consistent with previous studies using the same behavioral task (Miyazaki et al., 2014, Miyazaki et al., 2018) and confirmed that the administered optogenetic stimulation was sufficient to induce behavioral changes. These effects were similar for all 8 mice (Figure 3B), which suggests that the DRN was also effectively stimulated in the two mice in which we could not check the implantation site of the optic probes.

**Figure 3.**
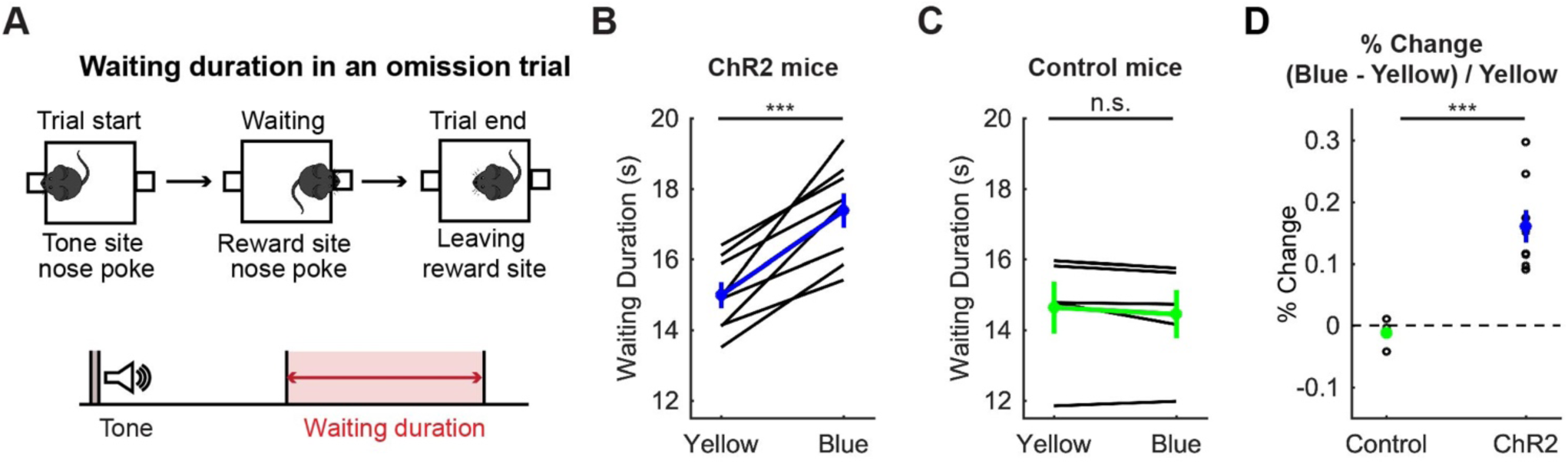
Activation of DRN 5-HT neurons prolonged stationary waiting for future rewards. (A) The definition of waiting duration. (B, C) Waiting duration in an omission trial in ChR2 (B, *n* = 8 mice) and control (C, *n* = 5 mice) mice. Blue and green dots indicate the mean across ChR2 and control mice, respectively. *** indicates p < 0.001 by paired t-test. (D) Change of waiting duration in blue light trials to yellow light trials in control (*n* = 5 mice) and ChR2 (n = 8 mice) mice. Green- and blue-filled circles indicate the mean across control (n = 5 mice) and ChR2 (n = 8 mice) mice respectively. *** indicates p < 0.001, unpaired t-test. n.s. indicates no significance (p > 0.05). Error bars represent the SEM in all graphs.

#### 3.2.2 Activation of DRN 5-HT neurons neither enhanced nor suppressed sustained motor actions

In order to examine whether optogenetic activation of DRN 5-HT neurons affects persistence in motor actions for future rewards, we analyzed the successful trial rate, the duration, the number of lever presses in omission trials, the time spent in abandoning an omission trial, and the action speed in the lever-pressing task.

##### 3.2.2.1 Successful trial rate

We first calculated the percentage of successfully rewarded trials in 8-, 16-, 32-, and 64-press trials. In 8-, 16-, and 32-press trials, mice successfully obtained rewards almost 100% of the time. In 64-press trials, the successful trial rate decreased, but was not significantly different between blue light and yellow light stimulation (t_7_ = 0.23, p = 0.83, paired t-test; Figure 4A).

**Figure 4.**
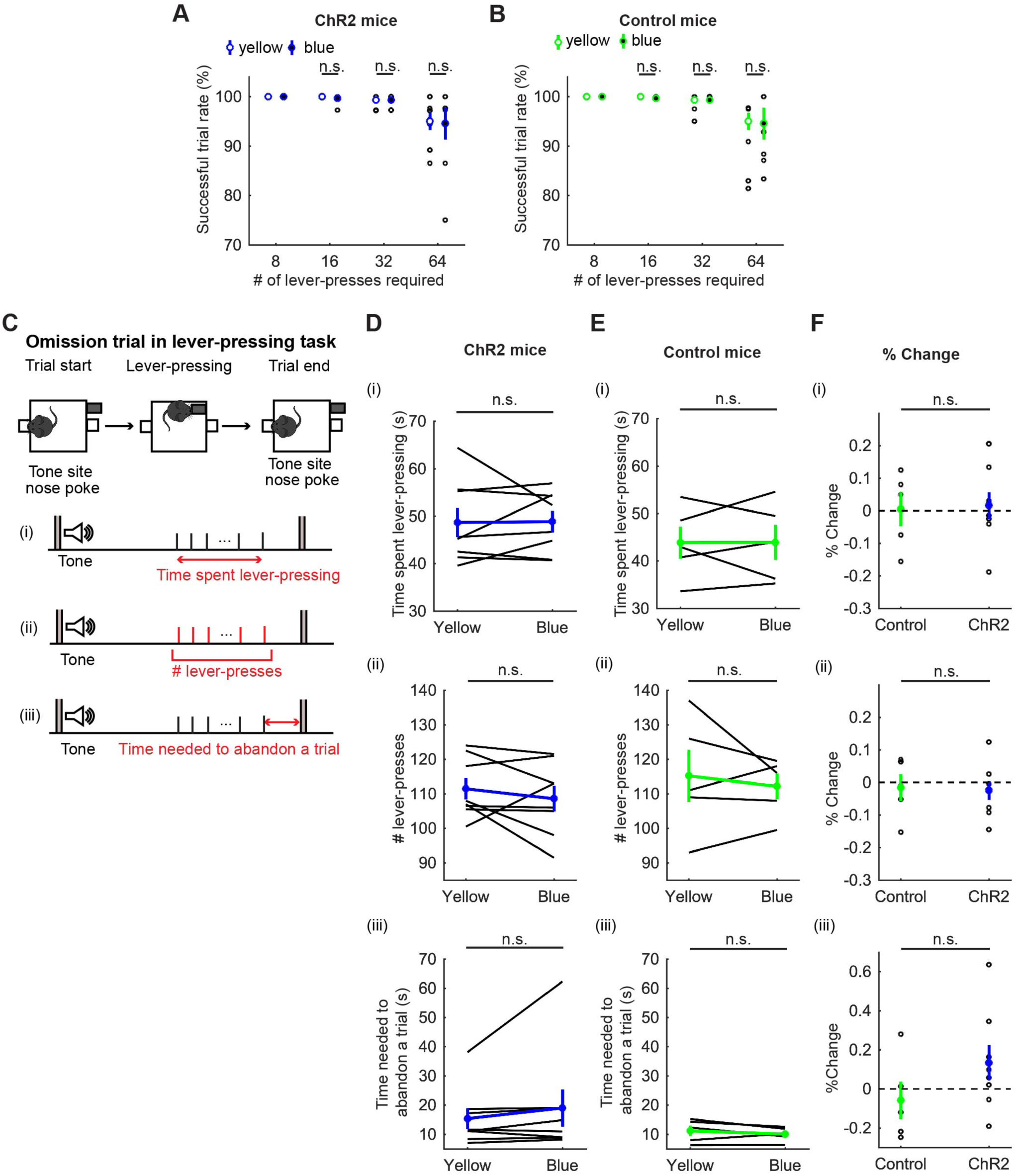
Activation of DRN 5-HT neurons did not change persistence in motor actions for future rewards. (A,B) Successful trial rates in ChR2 (A, *n* = 8 mice) and control (B, *n* = 5 mice) mice. Open and filled circles indicate the mean across blue and yellow light trials, respectively, in ChR2 (blue) and control (green) mice. n.s. indicates no significance (p > 0.05) by paired t-test. (C) The definition of behavioral measures for action persistence. (D, E) Behavioral parameters in ChR2 (D, *n* = 8 mice) and control (E, *n* = 5 mice) mice. Blue and green dots indicate the mean across ChR2 and control mice data, respectively. n.s. indicates no significance (p > 0.05) by paired t-test in D(i) and D(ii) and by Wilcoxon signed-rank test in D(iii). (F) Change of behavioral parameters in blue light trials to yellow light trials in control (*n* = 5 mice) and ChR2 (*n* = 8 mice) mice. Green- and blue-filled circles indicate the mean across control and ChR2 mice, respectively. n.s. indicates no significance (p > 0.05) by unpaired t-test. Error bars represent the SEM in all graphs.

##### 3.2.2.2 Time spent pressing the lever

To quantify how long mice could sustain actions for delayed rewards, we next measured the time spent pressing the lever and the elapsed time from the first lever-press to the last lever-press in omission trials (Figure 4C(i)). Interestingly, mice spent more than three times longer pressing the lever (48.68 ± 3.09 s with yellow light, Figure 4D(i)) than they spent in stationary waiting (15.0 ± 0.37 s, Figure 3B) for the same reward, showing that mice can tolerate longer delays while they are actively engaged in doing something, as opposed to waiting inactively. However, the time spent lever-pressing in an omission trial was not significantly different between trials with activation and those without (t_7_ = 0.083, p = 0.94, paired t-test; Figure 4D(i)). The change of duration in ChR2 was not significantly different from that in control mice (t_11_ = 0.16, p = 0.88, unpaired t-test; Figure 4D-F(i)).

##### 3.2.2.3 The number of lever-presses in omission trials

To quantify how persistently mice sustained motor actions for future rewards, we measured the number of lever-presses in omission trials (Figure 4D(ii)). The number of lever-presses in omission trials with optogenetic activation was not significantly different from that without activation (t_7_ = 0.93, p = 0.38, paired t-test; Figure 4D(ii)). The change in the number in ChR2 was not significantly different from that in control mice (t_11_ = 0.17, p = 0.87, unpaired t-test; Figure 4D-F(ii)).

##### 3.2.2.4 Time needed to abandon a trial

We next measured the time from the last lever-press to a nose poke in the tone site to abandon an omission trial, which could indicate how ambivalent mice were about abandoning the present trial (Figure 4C(iii)). In ChR2 mice, optogenetic activation did not significantly change the time spent to abandon a trial (z = 1.26, p = 0.23, Wilcoxon signed-rank test; Figure 4D(iii)) and the change in time was not significantly different from that in control mice (t_11_= 1.40, p = 0.19, unpaired t-test; Figure 4D-F(iii)).

##### 3.2.2.5 Action speed

In order to examine how optogenetic activation of DRN 5-HT neurons affects the speed of motor actions, we measured inter-press intervals (IPIs), the intervals between successive lever-presses. Behavioral observations indicated that mice usually pressed the lever continuously, but sometimes paused lever-pressing, such as resting, checking the reward site, or exploring, especially in omission trials. To be specific, most IPIs were < 5 s, but small numbers of IPIs were longer (Figure 5A). Therefore, we defined IPIs < 5 s as short IPIs, which represent continuous lever-pressing behavior, and IPIs ≥ 5 s as long IPIs, which mainly represent other behaviors, and examined the effect of the photoactivation on each type of IPI.

**Figure 5.**
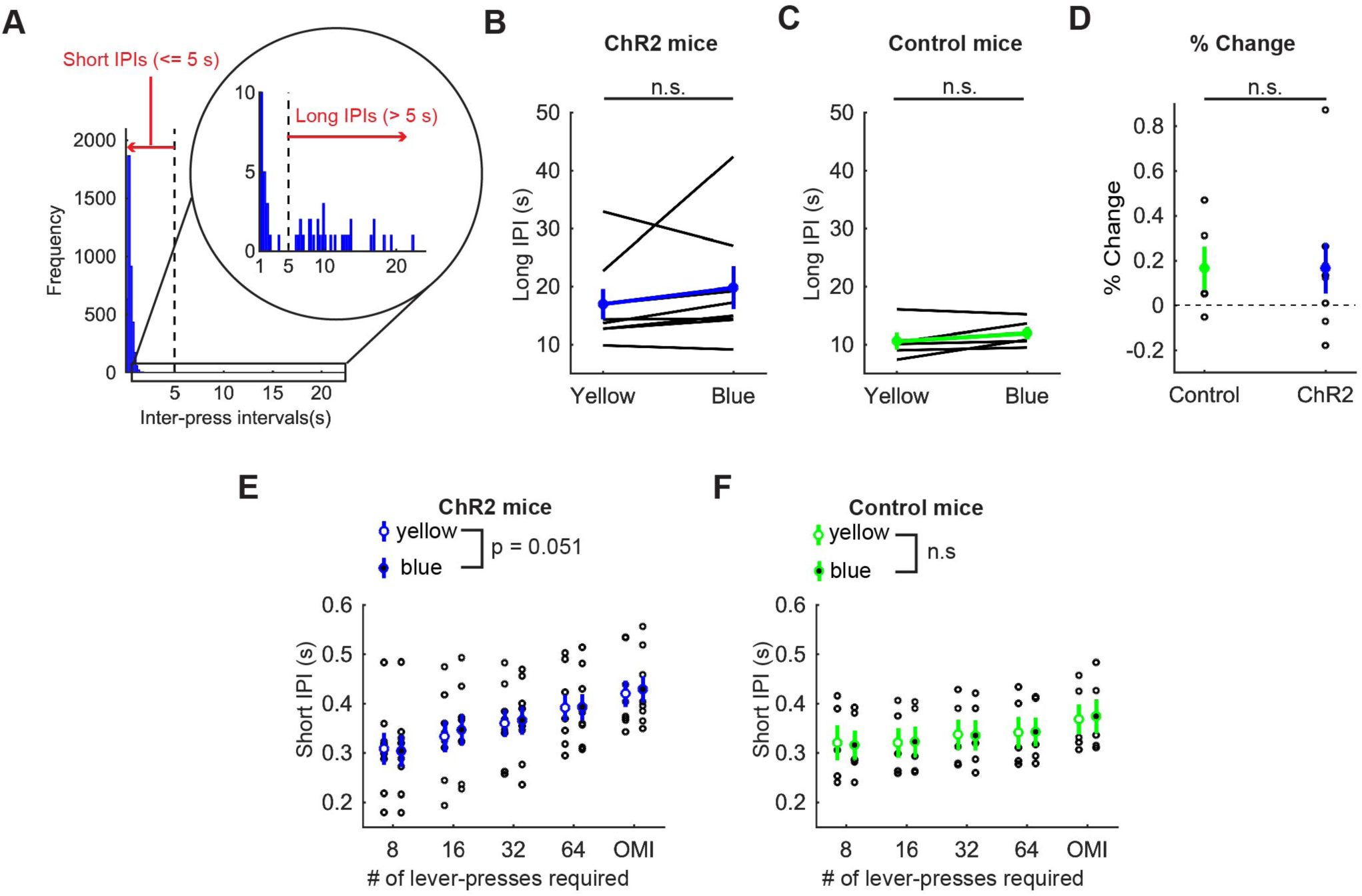
Activation of DRN 5-HT neurons did not change the speed of actions in the lever-pressing task. (A) The definition of long and short IPIs. (B,C) Long IPIs in omission trials in ChR2 (B, *n* = 8 mice) and control (C, *n* = 5 mice) mice. Blue and green dots indicate the means across ChR2 mice and control mice, respectively. n.s. indicates no significance (p > 0.05) by Wilcoxon signed-rank test in B and by paired t-test in C. (D) Change of long IPIs in blue light trials to yellow light trials in control (*n* = 5 mice) and ChR2 (*n* = 8 mice) mice. Green- and blue-filled circles indicate the means across control and ChR2 mice, respectively. n.s. indicates no significance (p > 0.05) by unpaired t-test. (E, F) Short IPIs in ChR2 (E, *n* = 8 mice) and control (F, *n* = 5 mice) mice. Open and filled circles indicate the mean across blue and yellow light trials respectively in ChR2 (blue) and control (green) mice. n.s. indicates no significance (p > 0.05) in the main effect of repeated measures ANOVA. Error bars represent the SEM in all graphs.

Long IPIs were analyzed only in omission trials, because they were rarely found in trials requiring 8 or more presses. In ChR2 mice, there was no significant difference in long IPIs between blue light and yellow light stimulation (z = 1.26, p = 0.23, Wilcoxon signed-rank test; Figure 5B). The change of long IPIs in ChR2 mice was not significantly different from that in control mice (t_11_ = 0.0027, p = 0.998, unpaired t-test; Figure 5B-D)).

For short IPIs, we first calculated the average short IPI in a trial and then the median across trials for each mouse. Those values were statistically tested using repeated measures ANOVA (Figure 5E for ChR2 and 5F for control mice). In ChR2 mice, there was a significant main effect of press conditions (five levels within-subject factors; 8-press, 16-press, 32-press, 64-press, and omission, *F*(4,28) = 25.64, *P* = 5.2 x 10^-9^) and a marginal main effect on the factor of stimulation conditions (yellow and blue, *F*(1,7) = 5.55, *P* = 0.051). However, there was no significant interaction effect (stimulation x press, *F*(4,28) = 1.86, *P* = 0.15). In control mice, there was a significant main effect of press conditions (*F*(4,16) = 42.62, *P* = 2.4 x 10^-8^), but no significant main effect of stimulation (*F*(1,4) = 0.16, *P* = 0.71). We also did not find a significant interaction effect (stimulation x press, *F*(4,16) = 0.29, *P* = 0.88).

##### 3.2.2.6 Premature reward checking

After repeated lever-presses, mice sometimes checked the reward site before reward delivery and then continued pressing the lever (58.66 ± 20.07 % of all omission trials). We define premature reward checking as a nose-poke at a reward site longer than 0.1s when the reward was unavailable.

We found that lever-pressing behavior was quite different before and after the first premature reward check. Short IPIs for each subject increased significantly after the first premature reward check (before: 0.35 ± 0.11 s; after: 0.75 ± 0.91 s; t_12_ = 2.76, p = 0.02, paired t-test). This suggests that mice focused more on lever-pressing behaviors before a premature reward check. Thus, we examined how activation of DRN 5-HT neurons affected action persistence and speed before a premature reward check. We analyzed behavioral measures using summary statistics as above, but we did not find a significant difference in the number of lever-presses or short IPIs (Figure S1).

#### 3.2.3 GLM analysis revealed that optogenetic activation slightly decreased action persistence and action speed

In order to dissociate the effect of optogenetic manipulation from other task-relevant factors like fatigue and satiety, as well as to consider the mixed effect of individual variability, we performed GLM analysis of the number and speed of lever presses before premature reward checks with regressors including optogenetic manipulation, elapsed time in a session, and the session number across the experimental period (see Section 2.9).

In ChR2 mice, the number of lever-presses slightly decreased as a result of optogenetic activation (Figure 6B Manipulation; t_611_ = 3.38, p = 0.00077), but not in control mice (Figure 6B Manipulation; t_390_ = 0.60, p = 0.55). Results of other fixed effect terms were similar across ChR2 and control groups. The number of lever-presses significantly decreased within a session (Figure 6B Elapsed time; ChR2 mice: t_611_ = 7.5, p = 2.2 x 10^-13^; Control mice: t_390_ = 8.22, p = 3.0 x 10^-15^) and significantly increased across sessions (**Figure** 6B # sessions; ChR2 mice: t_611_ = 5.3, p = 1.6 x 10^-7^; Control mice, t_390_ = 5.40, p = 1.1 x 10^-7^).

**Figure 6.**
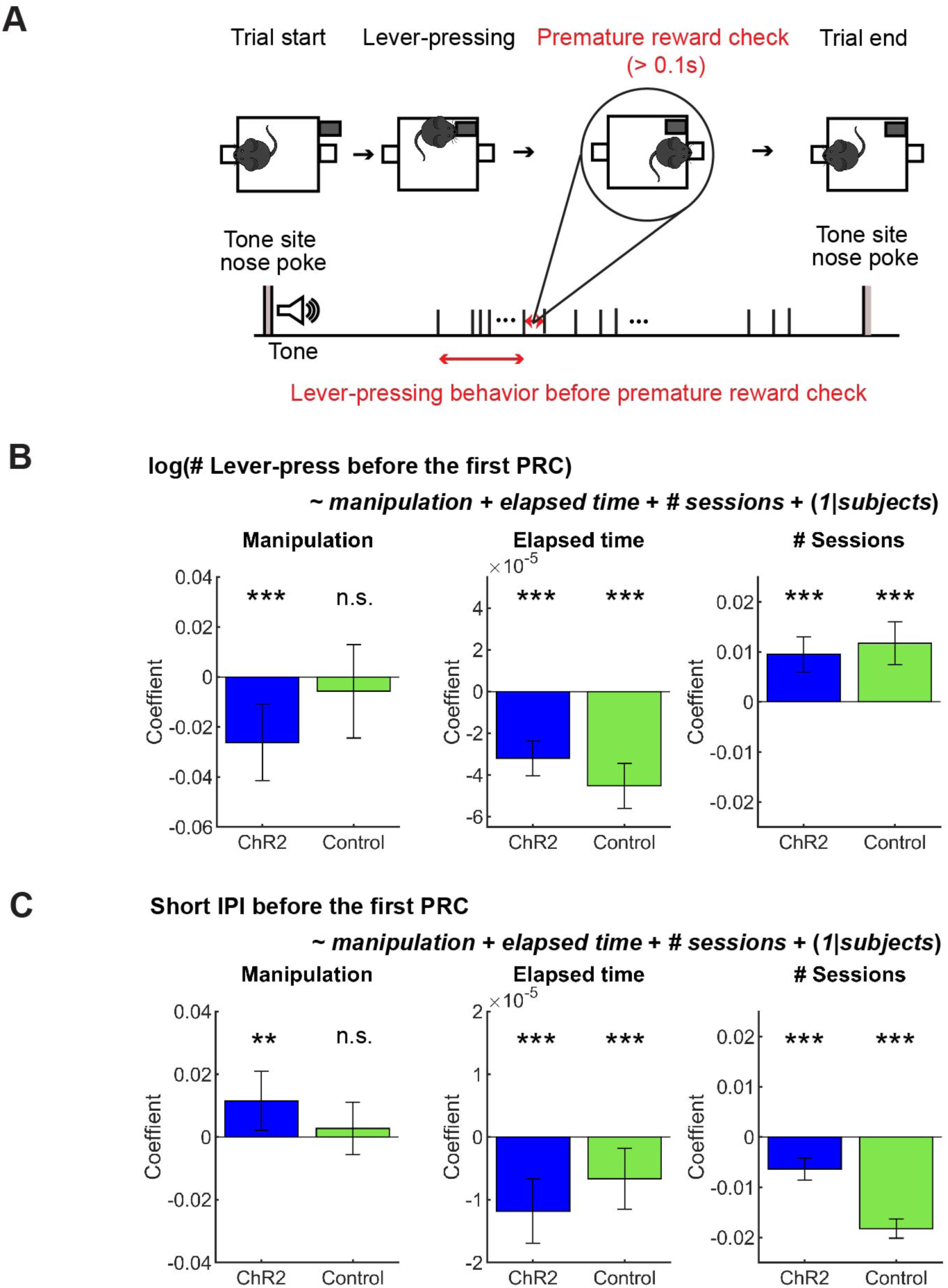
GLM analysis revealed a subtle effect of optogenetic activation on action persistence and speed. (A) The definition of premature reward check. (B) GLM model analysis of the number of lever-presses before the first premature check in Control (*n* = 5 mice) and ChR2 (*n* = 8 mice) mice. Blue and green bars indicate coefficients of optic stimulation (left), elapsed time (middle), and the number of sessions (right) in ChR2 and control mice, respectively. n.s. indicates no significance (p > 0.05) and *** (p > 0.001) indicates statistical significance by paired t-test. Error bars indicate 95% confidence intervals of the coefficients. (C) GLM model analysis of short IPIs before the first premature check in control (*n* = 5 mice) and ChR2 (*n* = 8 mice) mice. Blue and green bars indicate the coefficients of optic stimulation (left), elapsed time (middle), and the number of sessions (right) in ChR2 and control mice, respectively. n.s. indicates no significance (p > 0.05), and *** (p > 0.01) and *** (p > 0.001) indicates statistical significance by paired t-test. Error bars indicate 95 % confidence intervals of the coefficients.

Short IPIs were slightly increased by optogenetic activation in ChR2 mice, but not in control mice (Figure 6C Manipulation; ChR2 mice: t_611_ = 2.4, p = 0.017; t_390_ = 0.65, p = 0.52). Again, the results of other fixed-effect terms were consistent across the two groups. Short IPIs decreased within each session in both groups (Figure 6C Elapsed time; ChR2 mice: t_611_ = 4.50, p = 6.9 x 10^-6^); Control mice: t_390_ = 2.69, p = 0.0074) and decreased across multiple sessions (Figure 6C # sessions; ChR2 mice: t_611_ = 5.75, p = 1.3 x 10^-8^; Control mice: t_390_ = 18.61, p = 9.2 x 10^-56^).

In the results above, an interesting observation was that mice invested about three-fold more time for the same reward in the lever pressing task (Figure 4D) as in the stationary waiting task (Figure 3B), suggesting that lever pressing does not simply serve as a motor cost. While optogenetic activation of DRN 5-HT neurons clearly facilitated stationary waiting, its effect on lever presses was more subtle, and was detected only by focusing on the period before premature reward check and by subtracting the trends within and across sessions in GLMs.

### 3.3 Optogenetic inhibition of DRN 5-HT neurons

A subset of putative DRN 5-HT neurons increase their activity by behavioral activation, such as locomotion, changing direction, and approach/withdrawal behaviors (Ranade and Mainen, 2009). Given this observation, it is possible that lever-pressing behavior itself increases activity of DRN 5-HT neurons, such that optogenetic activation does not induce additional effects. Therefore, we examined the effect of optogenetic inhibition of DRN 5-HT neurons on action maintenance. In order to optogenetically inhibit DRN 5-HT neurons, continuous yellow light was applied from the onset of action until the end of the trial (Figure 1C).

#### 3.3.1 Inhibition of DRN 5-HT neurons shortened stationary waiting for future rewards

We first trained five ArchT and four control mice for the stationary waiting task and tested the effect of optogenetic inhibition of DRN 5-HT neurons. A previous study showed that chemical inhibition of DRN 5-HT neurons increased premature abandoning in a delayed reward task (Miyazaki et al., 2012b). Therefore, we predicted that optogenetic inhibition of DRN 5-HT neurons would decrease waiting duration in omission trials. As predicted, there was a significant decrease in waiting duration in omission trials during optogenetic inhibition (no light vs. yellow trials: t_4_ = 10.27, p = 0.00051, paired t-test; Figure 7B). The change in ArchT mice was significant compared to that in control mice (t_7_ = 4.07, p = 0.0047, unpaired t-test; Figure 7B-D). These results confirmed the effectiveness of the optogenetic inhibition protocol and also the causal relationship between decreased DRN 5-HT neural activity and impaired waiting for delayed rewards.

**Figure 7.**
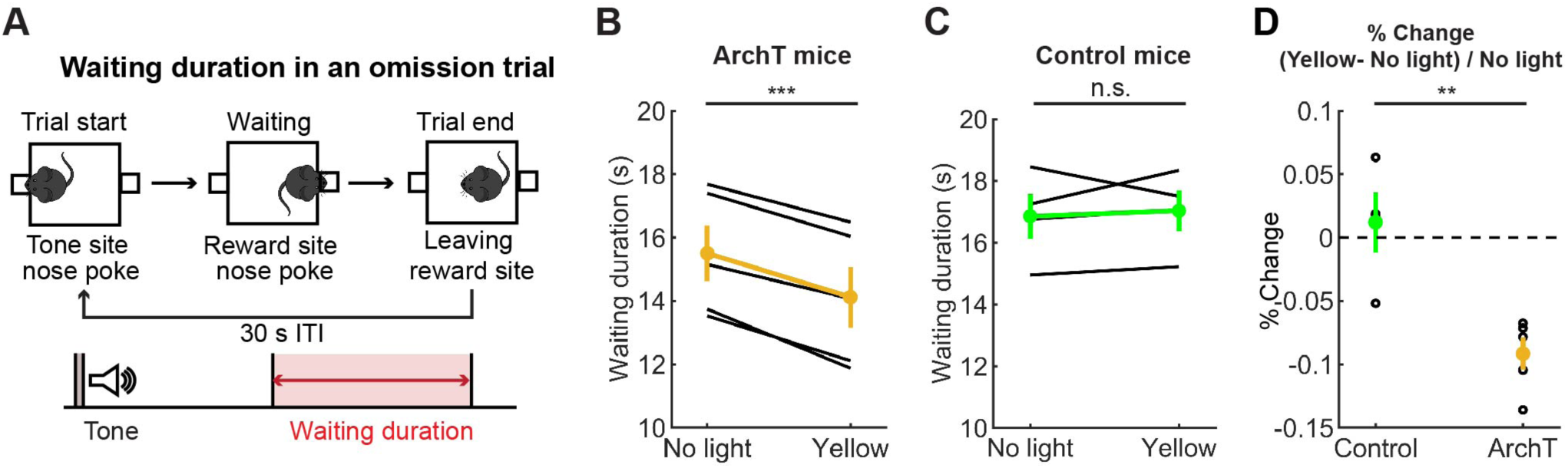
Inhibition of DRN 5-HT neurons shortened the stationary waiting period for future rewards. (A) The definition of waiting duration. (B, C) Waiting duration in an omission trial in ArchT (B, *n* = 5 mice) and control (C, *n* = 4 mice) mice. Yellow and green dots indicate the mean across ArchT and control mice, respectively. n.s. indicates no significance (p > 0.05) and *** indicates p < 0.01 by paired t-test. (D) Change of waiting duration in yellow light trials to no-light trials in control (*n* = 4 mice) and ArchT (*n* = 5 mice) mice. Green- and yellow-filled circles indicate the mean across control (*n* = 4 mice) and ArchT (*n* = 5 mice) mice, respectively. ** indicates p $<$ 0.01 by unpaired t-test. Error bars represent the SEM in all graphs.

#### 3.3.2 Inhibition of DRN 5-HT neurons neither enhanced nor suppressed sustained motor actions

In order to examine whether optogenetic inhibition of DRN 5-HT neurons affects sustained motor actions for future rewards, we analyzed the successful trial rate, the duration, the number of lever presses in omission trials, the time spent abandoning a trial, and the vigor in the lever-pressing task as in the optogenetic activation experiment.

##### 3.3.2.1 Successful trial rate

All mice were able to perform 8-, 16-, and 32-press trials without failure. In 64-press trials, the successful trial rate slightly decreased compared with other types of trials, but there was no significant difference between no-light and yellow-light trials (t_5_ = 0.24, p = 0.82, paired t-test; Figure 8A).

**Figure 8.**
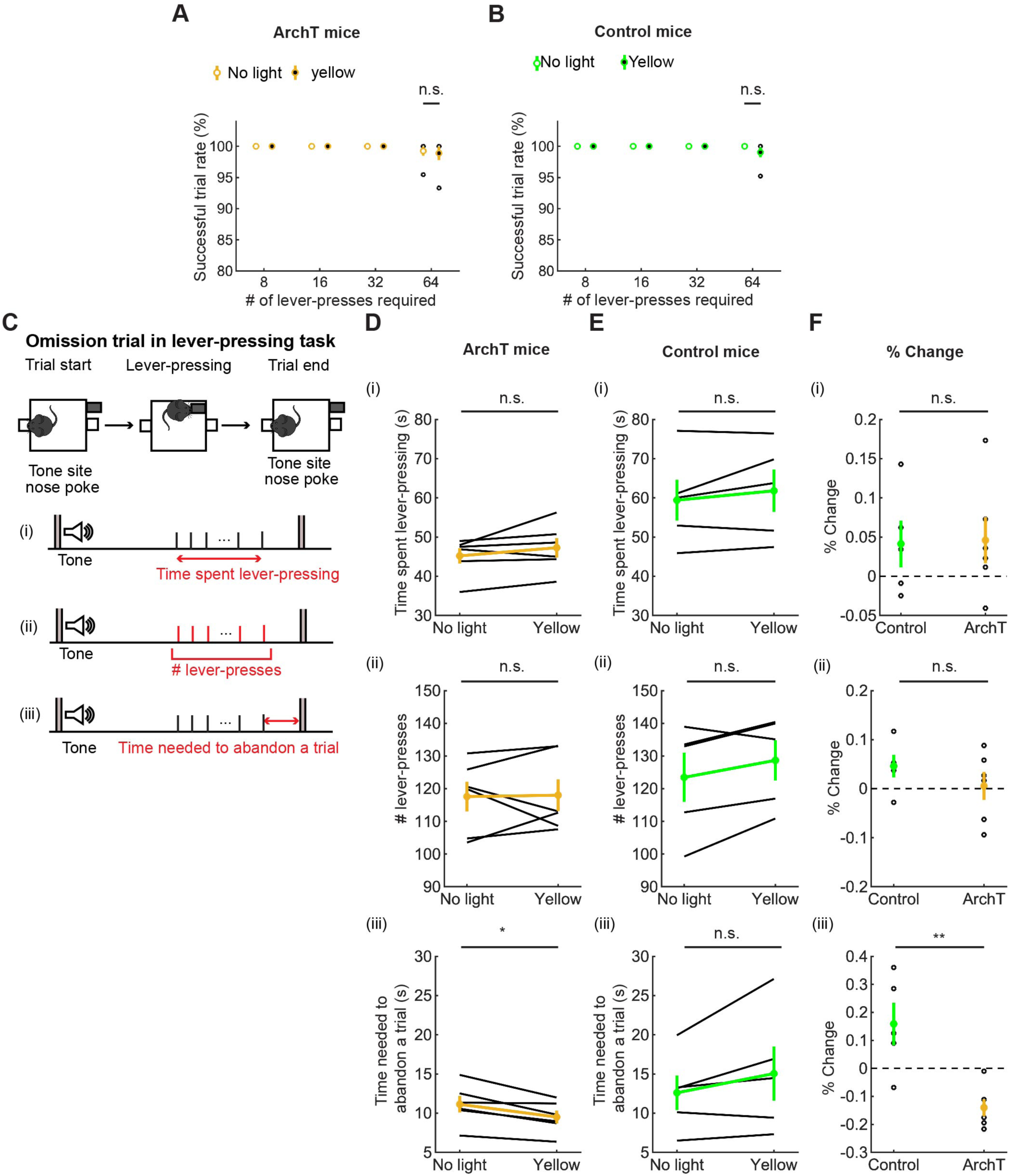
Inhibition of DRN 5-HT neurons neither enhanced nor suppressed sustained motor actions. (A,B) Successful trial rates in ArchT (A, *n* = 6 mice) and control (B, *n* = 5 mice) mice. Open and filled circles indicate the mean across no light and yellow light trials, respectively, in ArchT (yellow) and control (green) mice. n.s. indicates no significance (p $>$ 0.05) by paired t-test. (C) The definition of behavioral measures for action persistence. (D, E) Behavioral parameters in ArchT (D, *n* = 6 mice) and control (E, *n* = 5 mice) mice. Yellow and green dots indicate the mean across ArchT and control mice data, respectively. n.s. indicates no significance (p > 0.05) by Wilcoxon signed-rank test in D(i) and D(ii) and by paired t-test in other measures. (F) Change of behavioral parameters in yellow light trials to no-light trials in control (*n* = 5 mice) and ArchT (*n* = 6 mice) mice. Green- and yellow-filled circles indicate the means across control and ArchT mice, respectively. n.s. indicates no significance (p > 0.05) by unpaired t-test. Error bars represent the SEM in all graphs.

##### 3.3.2.2 Time spent pressing the lever

We measured the time that mice spent pressing the lever and the time from the first to the last lever-press, in omission trials. The time spent lever-pressing in omission trials (Figure 8C(i)) was much longer than that for stationary waiting in omission trials (Figure 7), as in the optogenetic activation experiment, but did not differ significantly between trials with and without optogenetic inhibition (W = 4.0, p = 0.22, Wilcoxon signed-rank test; Figure 8D(i)). The change of time in ArchT mice was not significantly different from that in control mice (t_9_ = 0.11, p = 0.91, unpaired t-test; Figure 8D-F(i)).

##### 3.3.2.3 The number of lever-presses in omission trials

We next measured the number of lever presses in omission trials. That number did not differ significantly between trials with and without optogenetic inhibition (W = 10, p = 1.0, Wilcoxon signed-rank test; Figure 8D(ii)). The change of the number in ArchT was not significantly different from that in control mice (t_9_ = 1.07, p = 0.31, unpaired t-test; Figure 8D-F(ii)).

##### 3.3.2.4 The time needed to abandon a trial

We also measured the time from the last lever-press to a nose poke in the tone site to abandon an omission trial. The duration significantly decreased as a result of optogenetic inhibition in ArchT mice (t_5_ = 3.69, p = 0.014, paired t-test; Figure 8D(iii)). On the other hand, the duration did not change in control mice (t_4_ = 1.77, p = 0.15, paired t-test; Figure 8E(iii)) and the change of time was significantly different in ArchT mice compared to control mice (t_9_ = 3.93, p = 0.0034, unpaired t-test; Figure 8F(iii)). Thinking together with decreased duration of waiting in the stationary waiting task, the specific effect of optogenetic inhibition on the time to abandon a trial suggests that serotonergic activity favorably modulates decisions based on time rather than motor actions.

##### 3.3.2.5 Action speed

To examine the effect of optogenetic inhibition on the speed of motor actions, we measured IPIs. As in the optogenetic activation experiment, we examined the effect of optogenetic inhibition on long and short IPIs (Figure 9A). Long IPIs in omission trials did not change significantly due to optogenetic inhibition (W = 7.0, p = 0.56, Wilcoxon signed-rank test; Figure 9B). The change of long IPI in ArchT was not significantly different from that of control mice (U = 21.0, p = 0.33, U test; Figure 9B-D)). We next analyzed data of ArchT and control mice with repeated measures ANOVA (Figure 9E for ArchT and 9F for control mice). In ArchT mice, no significant main effect was found due to stimulation conditions (main effects of stimulation: two levels within-subject factors; No light and yellow, *F*(1,5) = 0.28, *P* = 0.87) and press conditions (five levels within-subject factors; 8-press, 16-press, 32-press, 64-press, and omission, *F*(4,20) = 1.86, *P* = 0.16). There was no significant interaction effect (stimulation x press, *F*(4,20) = 2.53, *P* = 0.073). Similarly, in control mice, there was no significant main effect of stimulation (*F*(1,4) = 0.031 p = 0.87) and press (*F*(4,16) = 1.27, *P* = 0.32). There were no significant interaction effects (stimulation x press, *F*(4,16) = 0.37, *P* = 0.83).

**Figure 9.**
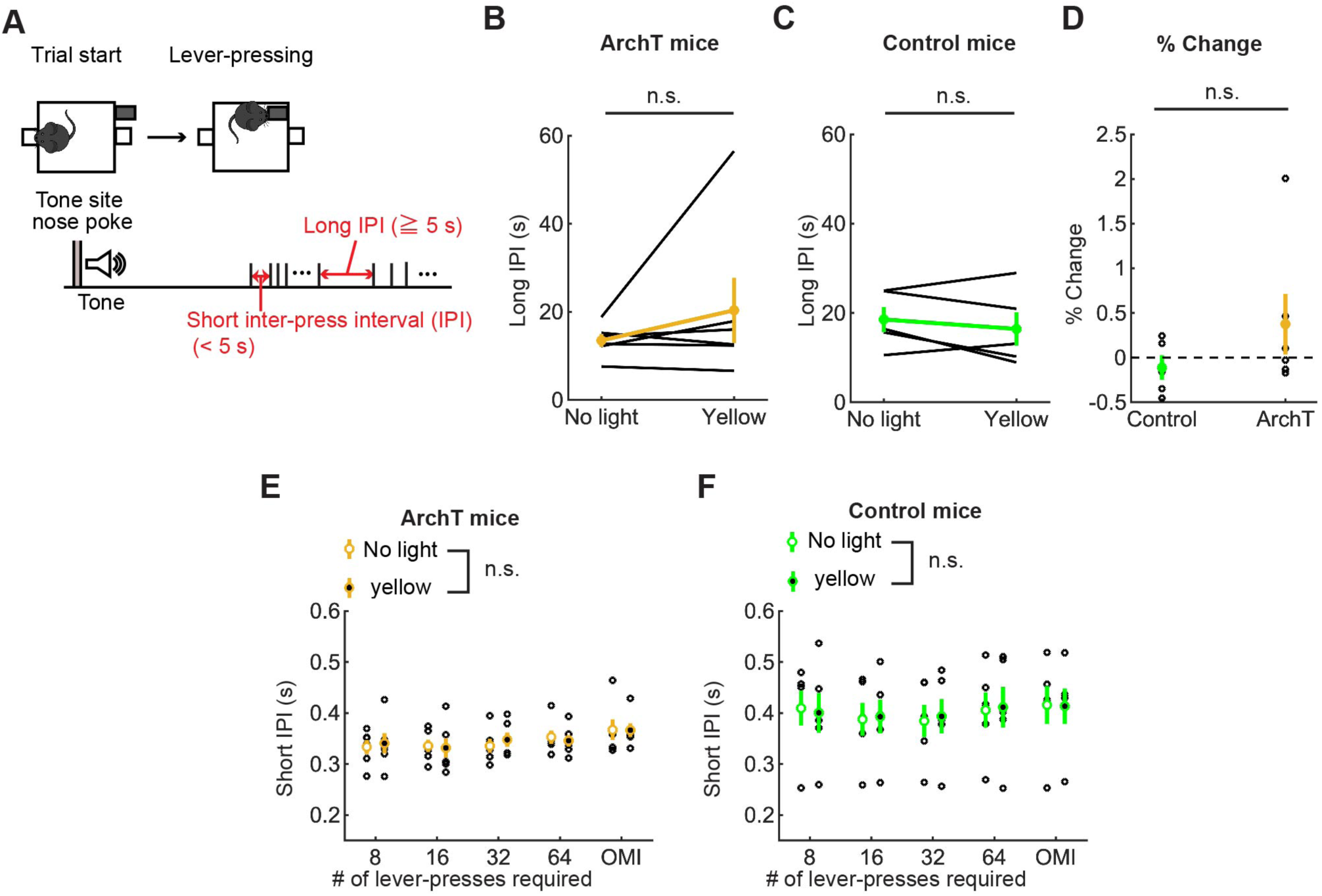
Inhibition of DRN 5-HT neurons did not change the speed of actions in the lever-pressing task. (A) Definitions of long and short IPIs. (B, C) Long IPIs in omission trials in ArchT (B, *n* = 6 mice) and control (C, *n* = 5 mice) mice. Yellow and green dots indicate the means across ArchT mice and control mice, respectively. n.s. indicates no significance (p > 0.05) by Wilcoxon signed-rank test. (D) Change of long IPIs in yellow light trials to no-light trials in control (*n* = 5 mice) and ArchT (*n* = 6 mice) mice. Green- and yellow-filled circles indicate the mean across control and ArchT mice, respectively. n.s. indicates no significance (p > 0.05) by Mann Whitney U-test. (E, F) Short IPIs in ArchT (E, *n* = 6 mice) and control (F, *n* = 5 mice) mice. Open and filled circles indicate the means across no-light and yellow light trials, respectively, in ArchT (yellow) and control (green) mice. n.s. indicates no significance in main effect of stimulation. Error bars represent the SEM in all graphs.

#### 3.3.3 Optogenetic inhibition did not change the number or speed of lever presses in GLM analysis

In order to further confirm the effect of optogenetic inhibition on action persistence and speed of motor actions, we examined the number of lever-presses and short IPIs before the first premature reward check in omission trials, as we did in optogenetic activation studies. In the analysis using summary statistics, optogenetic inhibition did not significantly change the number of lever-presses or of short IPIs (Figure S2).

In GLM analysis, the number of lever presses did not change as a result of optogenetic manipulation in either group (Figure 10A Manipulation; ArchT mice: t_227_ = −0.36, p = 0.72; Control mice: t_203_ = 1.30, p = 0.20). The results of other fixed effect terms were similar across ChR2 and control groups. The number of lever presses decreased in each session (Figure 10A Elapsed time; ArchT mice: t_227_ = 7.14, p = 1.2 x 10^-11^; Control mice: t_203_ = 8.45, p = 5.4 x 10^-15^) and increased across sessions (Figure 10A # Sessions; ArchT mice: t_227_ = 2.01, p = 0.046; Control mice, t_203_ = 2.77, p = 0.0061).

**Figure 10.**
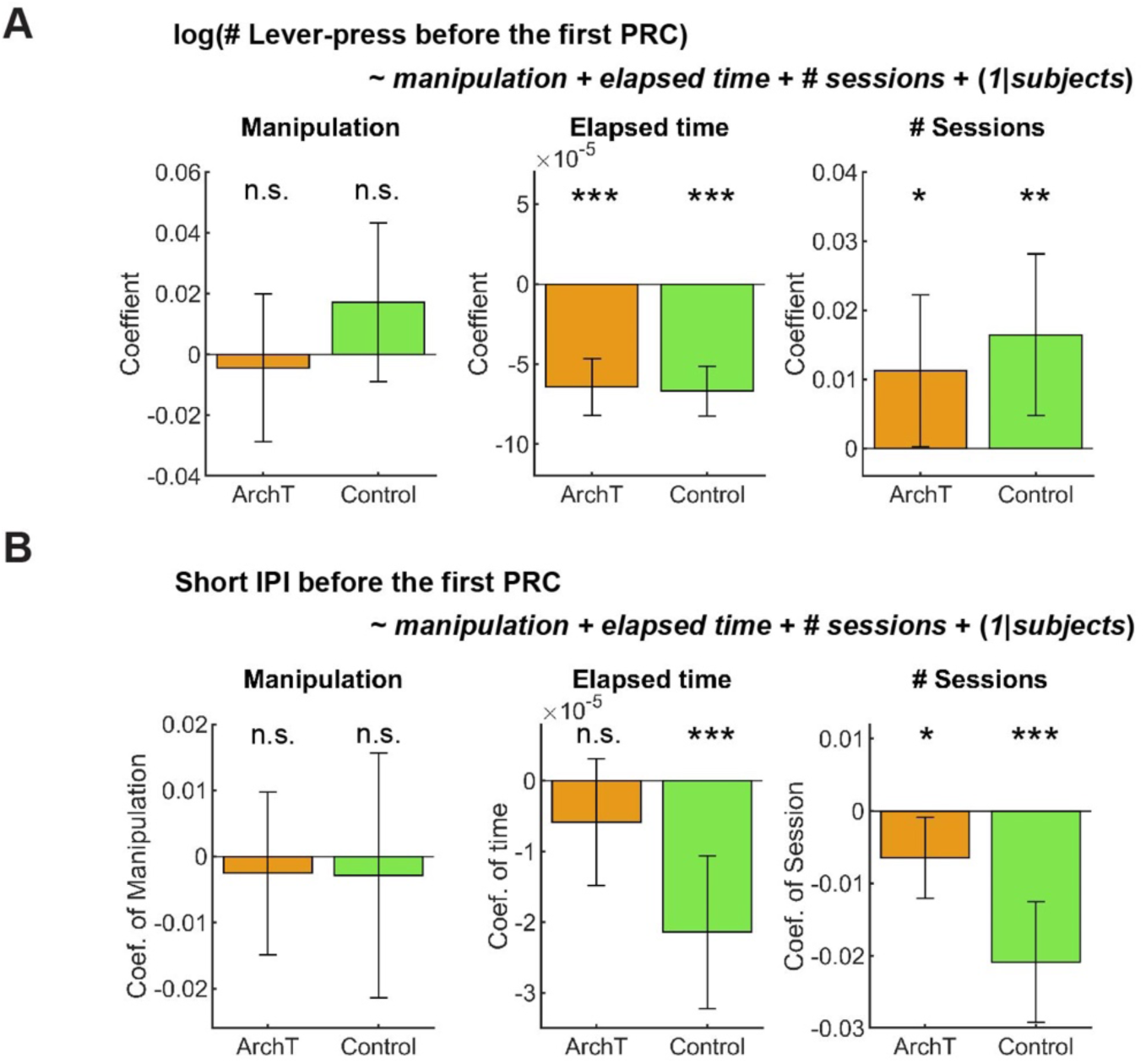
GLM analysis did not show the effect of optogenetic inhibition on active persistence or action vigor. (A) GLM analysis of the number of lever-presses before the first premature check in Control (*n* = 5 mice) and ArchT (*n* = 6 mice) mice. Orange and green bars indicate coefficients of optic manipulation (left), elapsed time (middle), and the number of sessions (right) in control and ArchT mice, respectively. n.s. indicates no significance (p > 0.05) and * (p > 0.05), ** (p > 0.01), and *** (p > 0.001) indicate statistical significance by paired t-test. Error bars indicate 95% confidence intervals of the coefficients. (B) GLM analysis of short IPIs before the first premature check in Control (*n* = 5 mice) and ArchT (*n* = 6 mice) mice. Orange and green bars indicate coefficients of optic manipulation (left), elapsed time (middle), and the number of sessions (right) in control and ArchT mice, respectively. n.s. indicates no significance (p > 0.05), and * (p > 0.05) and *** (p > 0.001) indicate statistical significance by paired t-test. Error bars indicate 95% confidence intervals of the coefficients.

Short IPIs did not show any significant change due to optogenetic manipulation (Figure 10B Manipulation; ArchT mice: t_227_ = 0.41, p = 0.68; Control mice: t_203_ = 0.31, p = 0.76). The results of other fixed effect terms were similar across both groups. In each session, short IPIs decreased in control mice (Figure 10B Elapsed time; t_203_ = 3.92, p = 0.00012), but not in ArchT mice (Figure 10B # Sessions; t_227_ = 1.3, p = 0.20). Across multiple sessions, short IPIs significantly decreased in both groups (ArchT mice: t_227_ = 2.3, p = 0.023; Control mice: t_203_ = 4.94, p = 1.6 x 10^-6^).

These results indicate that inhibition of DRN 5-HT neurons neither enhanced nor suppressed sustained motor actions. However, inhibition shortened duration of waiting in the stationary waiting task and the time to abandon a trial in the lever-pressing task, suggesting that DRN 5-HT neurons favorably modulate actions based on time, with limited effect on the cost of actions.

## 4 Discussion

In this study, we examined the role of DRN 5-HT neurons in sustaining motor actions for future rewards. We first showed that optogenetic activation of DRN 5-HT neurons prolonged waiting for future rewards, as in previous studies. On the other hand, the same stimulation neither enhanced nor suppressed the persistence and speed of motor actions in pooled analysis, i.e. summary statistics. In order to take into account trends within and across sessions and individual variability, we analyzed behavioral data using GLM, which revealed a subtle decrease in the number and speed of lever-presses due to optogenetic activation. To further investigate the causal relationship between activity of DRN 5-HT neurons and sustained motor actions, we optogenetically inhibited DRN 5-HT neurons during behavioral tasks. Optogenetic inhibition of DRN 5-HT neurons shortened the waiting duration, in opposition to its effect in the activation experiment. However, the same optogenetic inhibition did not induce changes in lever-pressing behavior either in summary statistics or GLM analysis, while it shortened the time to abandon a trial. Taken together, the present study revealed differential regulation of sustained motor actions and inactive waiting for future rewards with the main effect limited to waiting.

Based on the hypothesis that 5-HT controls temporal discounting of future rewards, we originally expected that optogenetic manipulation of DRN 5-HT neural activity modulates sustained motor actions for delayed rewards in much the same way as during stationary waiting. However, these experimental results suggest that the role of DRN 5-HT neurons in sustained motor actions cannot be fully explained by the discount factor hypothesis. Another possible interpretation of how activation of DRN 5-HT neurons prolongs waiting is based on the behavioral inhibition hypothesis, which suggests that increased 5-HT transmission shifts animal behavior toward inaction (Soubrie, 1986). If this hypothesis can account for 5-HT regulation of adaptive behaviors, it might be expected that activation of DRN 5-HT neurons would suppress lever-pressing behavior and vice versa. However, the present study showed a subtle decrease in lever-pressing behavior by optogentic activation and no effect on sustained motor actions by optogentic inhibition, suggesting that behavioral inhibition cannot explain the results of the present study.

In the present study, we found the differential optogenetic effect between stationary waiting and sustained motor actions. This may reflect distinctive neural substrates for delay-based (waiting) and effort-based (lever-pressing) motivated behaviors. A previous study showed that pharmacological reduction of systemic 5-HT level decreased the tendency of rats to favor immediate smaller rewards over delayed larger rewards in a delay-based choice task, but did not change their tendency to choose smaller rewards after climbing a low barrier rather than a larger reward after climbing a higher barrier in an effort-based choice task (Denk et al., 2005). Although sustained actions may be regulated differently than choice behaviors, our results are largely consistent with that behavioral study. Also, a recent fiber photometry study showed that DRN 5-HT neural activity is not increased while mice are pressing a lever to obtain rewards (Yoshida et al., 2019). Rather, the study showed that 5-HT neurons in the median raphe nucleus show increased neural activity during lever-pressing. How DRN 5-HT neurons regulate sustained actions differently has not been examined behaviorally, and our study adds behavioral evidence suggesting different neural substrates between sustained motor action and inactive waiting for future rewards.

A recent study showed that DRN 5-HT neurons are anatomically divided into at least two subtypes, cortex-projecting and subcortical projecting neurons, and that these subtypes show different responses to rewarding and punishing stimuli, and that they have different behavioral roles (Ren et al., 2018). Therefore, it is possible that DRN 5-HT projections to different brain regions have different behavioral functions. Supporting this notion, activation of serotonergic projections to the ventral tegmental area increased active nose-poking to receive optogenetic self-stimulation (Nagai et al., 2020), whereas pharmacological manipulation of 5-HT receptors has opposing effects on the vigor and persistence of actions, mainly through projections to the subsantia nigra or dorsomedial striatum (Bailey et al., 2018). In our study, we stimulated all DRN 5-HT neurons, which may have obscured the existence of functionally different DRN 5-HT projections on motor action for future rewards. Future experiments to stimulate axon terminals of DRN 5-HT projections to specific brain regions (Miyazaki et al., 2020) should further clarify the roles of DRN 5-HT neurons in sustained motor actions.

An interesting behavioral observation in the present study was that animals spent more time pressing a lever than continuing to nose poke in omission trials. That animals tolerate a longer delay to obtain the same reward during sustained active behavior than by stationary waiting is not compatible with the common notion that motor actions are costly. Previous psychological studies on self-control in humans showed that children who are engaged in behaviors to distract themselves, such as playing with toys or talking to others, can tolerate longer delays for rewards than those who did not (Mischel and Ebbesen, 1970, Mischel et al., 1972). From these studies, it was proposed that diverting their attention away from the temporal feature of the task by activating their behaviors reduces the effect of delay discounting (Metcalfe and Mischel, 1999). In the current study, mice may have been able to tolerate longer delays during the lever-pressing task due to actively sustained motor actions that shifted their attention away from the temporal feature of the task.

Another possible reason why animals can tolerate longer delays in lever-pressing is that lever-pressing behaviors are not as costly as inactive waiting. For example, voluntary running-wheel activity has properties of a positive reinforcer (Belke and Wagner, 2005). Based on the fact that mice were first trained to press a lever until it was withdrawn and reward was given to the food site, the tactile and auditory feedback of each lever-press would become conditioned reinforcer and would motivate mice to press longer. Previous studies showed that enhancement of serotonergic functions reduced the response to conditioned reinforcer (Guy and Fletcher, 2014, McDevitt et al., 2021) and voluntary running-wheel activity (Weber et al., 2009). If lever-pressing behavior has properties of both positive (conditioned) reinforcers and costly actions, serotonergic regulation of two behavioral processes, the decreased response to positive (conditioned) reinforcers and increased value of anticipated rewards over costly actions, might be confounding, which results in subtle or no effects by optogenetic manipulations.

Although the present study shows a subtle decrease in motor actions for future rewards by optogenetic activation, this result conflicts with the previous study, which showed that optogenetic activation of DRN 5-HT neurons promotes active persistence and increased response vigor in a task in which active nose poking was required to receive water rewards (Lottem et al., 2018) Also, the previous study differed from ours in the type of reward (water vs. food) and in actions associated with rewards (repeated nose pokes vs. lever-pressing). There are several possible reasons for the discrepancy.

The first is the difference in DRN 5-HT neural activity induced by optogenetic activation. In the previous study, ChR2(H134R) was expressed in SERT-Cre mice that received 25-Hz photostimulation. This induced strongly synchronized DRN 5-HT neural activity. Such strong neural activity was mainly observed at the time of reward acquisition (Li et al., 2016). A recent study using the same transgenic mouse line showed that 20-Hz photostimulation of DRN 5-HT projections to the ventral tegmental area induced dopamine release in the nucleus accumbens and also had a reinforcing effect (Wang et al., 2019). Several previous studies also showed that in the self-stimulation test, activation of DRN 5-HT neurons with 20-Hz photostimulation reinforced active nose poking to a port coupled with the stimulation (Liu et al., 2014, Nagai et al., 2020). Photostimulation used in the previous study may have affected active persistence and response vigor through the reward effect induced by dopamine release. On the other hand, in this study, we used ChR2(C128S), a step-type function opsin that induced less synchronized activity, i.e. approximately 6-Hz firing, and photostimulation to this ChR2 variant did not induce rewarding effects (Miyazaki et al., 2014), suggesting that dopamine release is not induced by the stimulation. One possibility is that moderate and strong activation of DRN 5-HT neurons interact with dopaminergic neurons differently, resulting in different effects on motivated behaviors. However, there are other possibilities discussed in previous studies, such as the difference in copy numbers of ChR2 expressing vectors/alleles (Nagai et al., 2020) or transgenic mouse lines (Cardozo Pinto et al., 2019). The two studies also have significant methodological differences to express ChR2 in DRN 5-HT neurons, i.e., injecting virus vector vs. crossing transgenic mouse lines. In order to clarify whether frequency stimulation or other factors cause these inconsistent results, it will be important to examine whether our stimulation in Tph2-ChR2(C128S) mouse lines induced behavioral effects observed in the previous study.

Another remarkable difference between our task and that of Lottem et al. (2018) is the cost of abandoning a trial. The amount of travel cost to move to the next trial critically modulates animal decisions about whether to stay in the current trial/patch. Previous studies found that time and response in the current trial increased as the travel cost increased (McCarthy et al., 1994, Vertechi et al., 2020). In Lottem et al. (2018), mice were required to travel a 30-cm passage to move to the next trial, which took approximately 3 s on average. On the other hand, in our task, the inter-trial intervals could function as a travel cost to start the next trial, which took at least 20 s. One possibility is that 5-HT neurons modulate active motor actions differently, depending on the magnitude of the travel cost. The travel cost can be important even for how 5-HT neurons modulate decisions based on the cost of time. In Lottem et al. (2018), optogenetic activation increased the time to leave a current trial. In the present study, we did not see a change in the time to abandon a trial in our optogenetic activation experiment, whereas optogenetic inhibition decreased the duration. This result supports the possibility that serotonergic regulation of sustained actions is dependent on future costs of travel time. This possibility can be examined by testing optogenetic stimulation of 5-HT neurons in lever-pressing behaviors with different lengths of inter-trial intervals.

In conclusion, we showed that manipulation of DRN 5-HT activity has different effects on sustained motor action and stationary waiting for future rewards. Patience to wait and persistence to act for delayed reward acquisition are regulated differently by DRN 5-HT neurons. Advanced optical imaging from genetically tagged 5-HT neurons such as fiber photometry and endoscopic microscopy will enable us to identify different neural substrates in persistence to act and patience to wait.

## Ethics Statement

The animal study was reviewed and approved by the Okinawa Institute of Science and Technology Experimental Animal Committee.

## Author Contributions

MT, KWM, KM, and KD designed the experiments. MT, JC, SOS, MN, TM conducted the behavioral experiments and analyzed the data. MT, JC and AC performed histological experiment and acquisition of histological images. AY and KFT generated Tph2-ChR2(C128S)-EYFP and Tph2-ArchT-EGFP bi-transgenic mice. MT wrote the original draft of the manuscript, and JC, KWM, KM, and KD edited the manuscript. All authors reviewed the manuscript.

## Acknowledgments

We thank Prof. Jeff Wickens, Julie Chouinard, Yumiko Akamine, Mao-Ting Hsu, Ai Takahashi, and Yuichi Morohashi for helpful guidance and advice on the histological experiments. We also thank Paolo Barzaghi in the OIST imaging section for help with cell counting work.

## Data Availability Statement

Data from the experiments presented in this study are available from the corresponding author to the responsible request.

