## Supplementary Figures for "The differential effect of optogenetic serotonergic manipulation on sustained motor actions and stationary waiting for future rewards in mice"

**Table S1. Statistical details for optogenetic activation study**

| Fig. | Sample size | Mean and SEM | Statistical test | Values |
|---|---|---|---|---|
| 3B | 8 ChR2 mice | Yellow: 14.99 ± 0.37 s<br>Blue: 17.39 ± 0.49 s | paired t-test | $t_7$ = 6.31, p = 0.00040 |
| 3C | 5 control mice | Yellow: 14.63 ± 0.74 s<br>Blue: 14.45 ± 0.68 s | paired t-test | $t_4$ = 1.50, p = 0.21 |
| 3D | 8 ChR2 mice<br>5 control mice | ChR2: 0.16 ± 0.027<br>Control: -0.012 ± 0.0087 | unpaired t-test | $t_{11}$ = 4.98, p = 0.00042 |
| 4A | 8 ChR2 mice | 64 press yellow:<br>95.02 ± 1.73%<br>64 press blue:<br>94.55 ± 3.23% | paired t-test | $t_7$ = 0.23, p = 0.83 |
| 4B | 5 control mice | 64 press yellow:<br>90.08 ± 3.46%<br>64 press blue:<br>90.33 ± 2.86% | paired t-test | $t_4$ = 0.073, p = 0.95 |
| 4D(i) | 8 ChR2 mice | Yellow: 48.68 ± 3.09 s<br>Blue: 48.86 ± 2.27 s | paired t-test | $t_7$ = 0.083, p = 0.94 |
| 4D(ii) | 8 ChR2 mice | Yellow: 111.50 ± 3.09<br>Blue: 108.63 ± 3.74 | paired t-test | $t_7$ = 0.93, p = 0.38 |
| 4D(iii) | 8 ChR2 mice | Yellow: 15.41 ± 3.53 s<br>Blue: 19.03 ± 6.37 s | signed-rank test | z = 1.26, p = 0.23 |
| 4E(i) | 5 control mice | Yellow: 43.86 ± 3.39 s<br>Blue: 43.92 ± 3.73 s | paired t-test | $t_4$ = 0.026, p = 0.98 |
| 4E(ii) | 5 control mice | Yellow: 115.2 ± 7.55 Blue:<br>112.2 ± 3.74 | paired t-test | $t_4$ = 0.58, p = 0.59 |
| 4E(iii) | 5 control mice | Yellow: 11.24 ± 1.75 s<br>Blue: 10.09 ± 1.10 s | paired t-test | $t_4$ = 1.11, p = 0.33 |
| 4F(i) | 8 ChR2 mice<br>5 control mice | ChR2: 0.016 ± 0.042<br>Control: 0.0048 ± 0.0052 | unpaired t-test | $t_{11}$ = 0.16, p = 0.88 |
| 4F(ii) | 8 ChR2 mice<br>5 control mice | ChR2: -0.024 ± 0.029<br>Control: -0.016 ± 0.041 | unpaired t-test | $t_{11}$ = 0.17, p = 0.87 |
| 4F(iii) | 8 ChR2 mice<br>5 control mice | ChR2:0.13 ± 0.090<br>Control: -0.059 ± 0.096 | unpaired t-test | $t_{11}$ = 1.40, p = 0.19 |
| 5B | 8 ChR2 mice | Yellow: 16.94 ± 2.65 s<br>Blue: 19.80 ± 3.70 s | signed-rank test | z = 1.26, p = 0.23 |

| | | | | |
|---|---|---|---|---|
| 5C | 5 control mice | Yellow: 10.56 ± 1.47 s<br>Blue: 11.92 ± 1.06 s | paired t-test | $t_4 = 1.61$, $p = 0.18$ |
| 5D | 8 ChR2 mice<br>5 control mice | ChR2: 0.17 ± 0.11<br>Control: 0.17 ± 0.096 | unpaired t-test | $t_{11} = 0.0027$, $p = 0.998$ |

**Table S2. Details of repeated measures ANOVA analysis of short IPIs in the optogenetic activation studies**

| Sample size | Factors | Values |
|---|---|---|
| 8 ChR2 mice | Stimulation (2 level, within)<br>Press (5 level, within) | Stimulation: $F(1,7) = 5.55$, $P = 0.051$<br>Press: $F(4,28) = 25.64$, $P = 5.2 \times 10^{-9}$<br>Interaction: $F(4,28) = 1.86$, $P = 0.15$ |
| 5 control mice | Stimulation (2 level, within)<br>Press (5 level, within) | Stimulation: $F(1,4) = 0.16$, $P = 0.71$<br>Press: $F(4,16) = 42.62$, $P = 2.4 \times 10^{-8}$<br>Interaction: $F(4,16) = 0.29$, $P = 0.88$ |

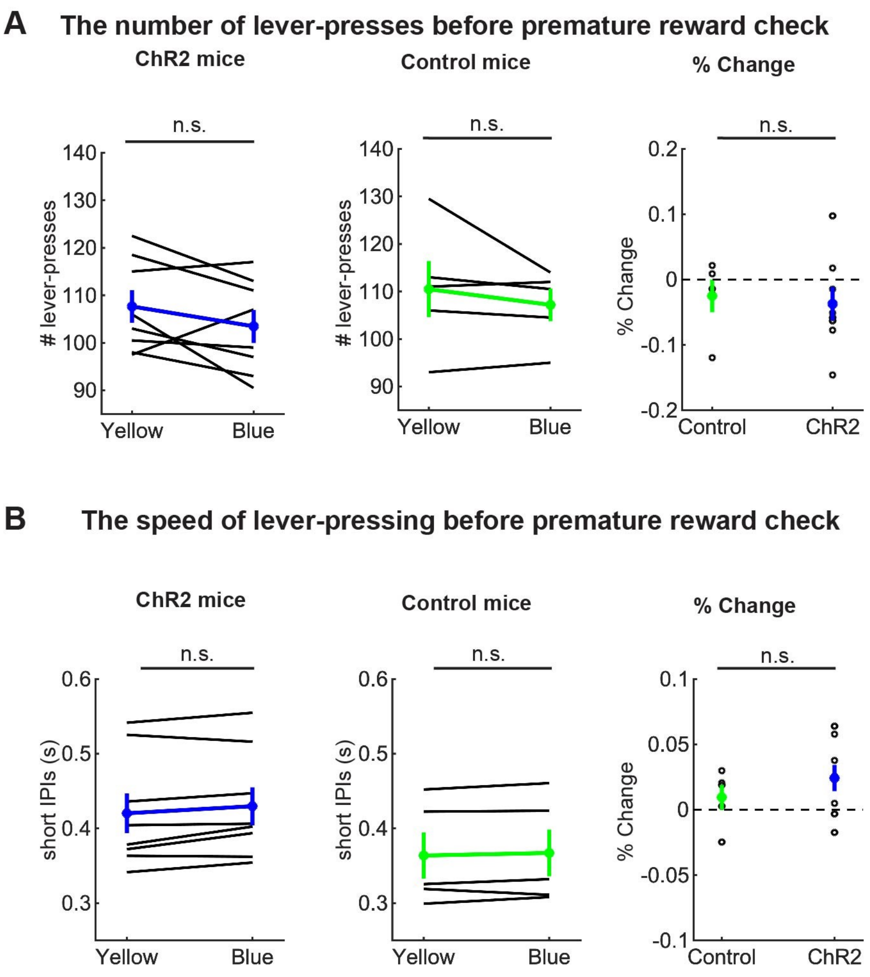


**Figure S1. Optogenetic inhibition did not change the summary statistics of action persistence and speed before premature reward check in summary statistics** (A) The number of lever-presses before premature reward check in omission trials in ChR2 (Left, *n* = 8 mice) and control (Middle, *n* = 5 mice) mice. Blue and green dots indicate the means across ChR2 and control mice data, respectively. Right panel indicates change of the number of lever-presses in blue light trials to yellow light trials in control (*n* = 5 mice) and ChR2 (*n* = 8 mice) mice. Green and blue-filled circles indicate the means across control and ChR2 mice, respectively. n.s. indicates no significance ($p > 0.05$). (B) Short IPIs before premature reward check in omission trials in ChR2 (Left, *n* = 8 mice) and control (Middle, *n* = 5 mice) mice. Blue and green dots indicate the means across ChR2 and control mice data, respectively. Right panel indicates the change of the short IPIs in blue light trials to yellow light trials in control (*n* = 5 mice) and ChR2 (*n* = 8 mice) mice. Green- and blue-filled circles indicate the means across control and ChR2 mice, respectively. n.s. indicates no significance ($p > 0.05$). See Table S3 for the detail of statistical tests. Error bars represent the SEM in all graphs.

**Table S3. Statistical details for Figure S1**

| Fig. | Sample size | Mean and SEM | Statistical test | Values |
|---|---|---|---|---|
| S1A | 8 ChR2 mice | Yellow: 107.63 ± 3.44<br>Blue: 103.44 ± 3.49 | paired t-test | $t_7 = 1.56$, p = 0.16 |
| S1A | 5 control mice | Yellow: 110.50 ± 5.89<br>Blue: 107.20 ± 3.44 | paired t-test | $t_4 = 1.05$, p = 0.21 |
| S1A | 8 ChR2 mice<br>5 control mice | ChR2: -0.037 ± 0.026<br>Control: -0.025 ± 0.025 | unpaired t-test | $t_{11} = 0.31$, p = 0.76 |
| S1B | 8 ChR2 mice | Yellow: 0.42 ± 0.027 s<br>Blue: 0.43 ± 0.026 s | paired t-test | $t_7 = 2.32$, p = 0.053 |
| S1B | 5 control mice | Yellow: 0.36 ± 0.031 s<br>Blue: 0.37 ± 0.032 s | paired t-test | $t_4 = 1.10$, p = 0.33 |
| S1B | 8 ChR2 mice<br>5 control mice | ChR2: 0.024 ± 0.010<br>Control: 0.0095 ± 0.0096 | paired t-test | $t_7 = 0.99$, p = 0.35 |

**Table S4. GLM analysis of the number of lever-press in the optogenetic activation study**

| Group | Effect | Estimate | 95% CI lower / upper | t-stats | p-values |
|---|---|---|---|---|---|
| ChR2 | Intercept | 4.70 | 4.66 / 4.75 | $t_{611} = 197.18$ | 0 |
| | Manipulation | -0.026 | -0.041 / -0.011 | $t_{611} = 3.38$ | 0.00077 |
| | Elapsed time | $-3.2\times10^{-5}$ | $-4.0\times10^{-5}$ / $-2.4\times10^{-5}$ | $t_{611} = 7.5$ | $2.2\times10^{-13}$ |
| | Sessions | 0.0095 | 0.0060 / 0.0130 | $t_{611} = 5.3$ | $1.6\times10^{-7}$ |
| Control | Intercept | 4.73 | 4.68 / 4.78 | $t_{390} = 169.82$ | 0 |
| | Manipulation | -0.0057 | -0.024 / 0.013 | $t_{390} = 0.60$ | 0.55 |
| | Elapsed time | $-4.5\times10^{-5}$ | $-5.6\times10^{-5}$ / $-3.4\times10^{-5}$ | $t_{390} = 8.22$ | $3.0\times10^{-15}$ |
| | Sessions | 0.012 | 0.0074 / 0.016 | $t_{390} = 5.40$ | $1.1\times10^{-7}$ |

**Table S5. GLM analysis of short IPIs in the optogenetic activation studies**

| Group | Effect | Estimate | 95% CI lower / upper | t-stats | p-values |
|---|---|---|---|---|---|
| ChR2 | Intercept | 0.470 | 0.42 / 0.52 | $t_{611} = 19.34$ | $2.4\times10^{-65}$ |
| | Manipulation | 0.012 | 0.0021 / 0.021 | $t_{611} = 2.4$ | 0.017 |
| | Elapsed time | $-1.2\times10^{-5}$ | $-1.7\times10^{-5}$ / $-6.7\times10^{-6}$ | $t_{611} = 4.5$ | $6.9\times10^{-6}$ |
| | Sessions | -0.0064 | -0.0085 / -0.0042 | $t_{611} = 5.75$ | $1.3\times10^{-8}$ |
| Control | Intercept | 0.46 | 0.40 / 0.52 | $t_{390} = 15.80$ | $9.0\times10^{-44}$ |
| | Manipulation | 0.0028 | -0.0056 / 0.011 | $t_{390} = 0.65$ | 0.52 |
| | Elapsed time | $-6.6\times10^{-6}$ | $-1.1\times10^{-5}$ / $-1.8\times10^{-6}$ | $t_{390} = 2.69$ | 0.0074 |
| | Sessions | -0.018 | -0.020 / -0.016 | $t_{390} = 18.61$ | $9.2\times10^{-56}$ |

**Table S6. Statistical details of the optogenetic inhibition studies**

| Fig | Sample size | Mean and SEM | Statistical test | Values |
|---|---|---|---|---|
| 7B | 5 ArchT mice | No light: 15.50 ± 0.88 s<br>Yellow: 14.11 ± 0.96 s | paired t-test | $t_4$ = 10.27, p = 0.00051 |
| 7C | 4 control mice | No light: 16.86 ± 0.73 s<br>Yellow: 17.04 ± 0.66 s | paired t-test | $t_3$ = 0.43, p = 0.70 |
| 7D | 5 ArchT mice<br>4 control mice | ArchT: -0.092 ± 0.013<br>Control: 0.012 ± 0.024 | unpaired t-test | $t_7$ = 4.07, p = 0.0047 |
| 8A | 6 ArchT mice | 64 press no light:<br>99.24 ± 0.76%<br>64 press yellow:<br>98.89 ± 1.11% | paired t-test | $t_5$ = 0.24, p = 0.82 |
| 8B | 5 control mice | 64 press no-light:<br>100 ± 0%<br>64 press yellow:<br>99.05 ± 0.95% | paired t-test | $t_4$ = 1.0, p = 0.37 |
| 8D(i) | 6 ArchT mice | No light: 45.19 ± 1.97 s<br>Yellow: 47.25 ± 2.47 s | signed-rank test | W = 4.0, p = 0.22 |
| 8D(ii) | 6 ArchT mice | No light: 117.57 ± 4.54<br>Yellow: 118.01 ± 4.85 | signed-rank test | W = 10, p = 1.0 |
| 8D(iii) | 6 ArchT mice | No light: 11.14 ± 1.05 s<br>Yellow: 9.51 ± 0.82 s | paired t-test | $t_5$ = 3.69, p = 0.014 |
| 8E(i) | 5 control mice | No light: 59.41 ± 5.20 s<br>Yellow: 61.82 ± 5.44 s | paired t-test | $t_4$ = 1.33, p = 0.25 |
| 8E(ii) | 5 control mice | No light: 123.48 ± 7.53<br>Yellow: 128.65 ± 6.17 | paired t-test | $t_4$ = 2.03, p = 0.11 |
| 8E(iii) | 5 control mice | No light: 12.60 ± 2.21 s<br>Yellow: 15.05 ± 3.47 s | paired t-test | $t_4$ = 1.77, p = 0.15 |
| 8F(i) | 6 ArchT mice<br>5 control mice | ArchT: 0.046 ± 0.030<br>Control: 0.041 ± 0.030 | unpaired t-test | $t_9$ = 0.11, p = 0.91 |
| 8F(ii) | 6 ArchT mice<br>5 control mice | ArchT: 0.0056 ± 0.029<br>Control: 0.046 ± 0.023 | unpaired t-test | $t_9$ = 1.07, p = 0.31 |
| 8F(iii) | 6 ArchT mice<br>5 control mice | ArchT: -0.14 ± 0.031<br>Control: 0.16 ± 0.076 | unpaired t-test | $t_9$ = 3.93, p = 0.0034 |
| 9B | 6 ArchT mice | No light: 13.50 ± 1.52 s<br>Yellow: 20.31 ± 7.40 s | signed-rank test | W = 7.0, p = 0.56 |

| | | | | |
|---|---|---|---|---|
| 9C | 5 control mice | No light: 18.50 ± 2.82 s<br>Yellow: 16.42 ± 3.76 s | paired t-test | $t_4 = 0.92$, p = 0.41 |
| 9D | 6 ArchT mice<br>5 control mice | ArchT: 0.37 ± 0.34<br>Control: -0.11 ± 0.14 | U test | $U = 21.0$, p = 0.33 |

**Table S7. Details of repeated measures ANOVA analysis of short IPIs in the optogenetic inhibition studies**

| Sample size | Factors | Values |
|---|---|---|
| 6 ArchT mice | Stimulation (2 level, within)<br>Press (5 level, within) | Stimulation: $F(1,5) = 0.28$, $P = 0.87$<br>Press: $F(4,20) = 1.86$, $P = 0.16$<br>Interaction: $F(4,20) = 2.53$, $P = 0.073$ |
| 5 control mice | Stimulation (2 level, within)<br>Press (5 level, within) | Stimulation: $F(1,4) = 0.031$, $P = 0.87$<br>Press: $F(4,16) = 1.27$, $P = 0.32$<br>Interaction: $F(4,16) = 0.37$, $P = 0.83$ |

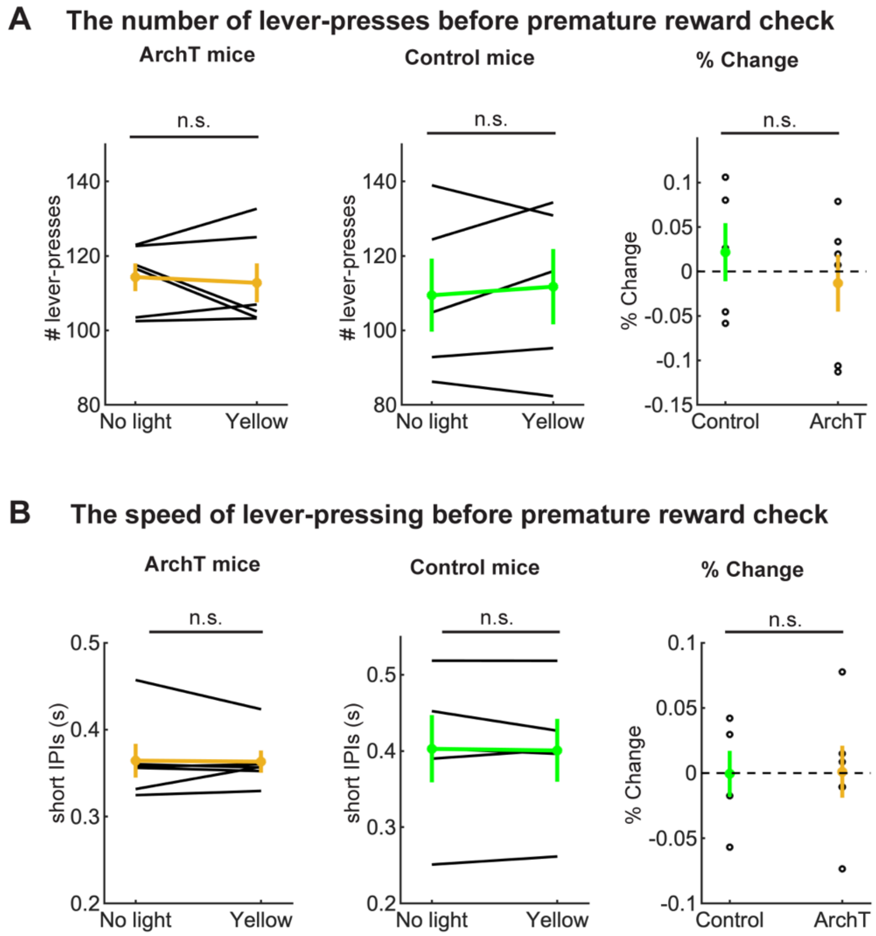


**Figure S2. Optogenetic inhibition did not change the summary statistics of action persistence and speed before premature reward check.** (A) The number of lever-presses before premature reward check in omission trials in ArchT (Left, $n = 6$ mice) and control (Middle, $n = 5$ mice) mice. Yellow and green dots indicate the means across ArchT and control mice, respectively. Right panel indicates change of the number of lever-presses in yellow light trials to no light trials in control ($n = 5$ mice) and ArchT ($n = 6$ mice) mice. Green- and yellow-filled circles indicate the means across control and ArchT mice, respectively. n.s. indicates no significance ($p > 0.05$). (B) Short IPIs before premature reward check in omission trials in ArchT (Left, $n = 6$ mice) and control (Middle, $n = 5$ mice) mice. Yellow and green dots indicate the means across ArchT and control mice, respectively. Right panel indicates the change of the short IPIs in yellow light trials to no light trials in control ($n = 5$ mice) and ArchT ($n = 6$ mice) mice. Green- and yellow-filled circles indicate the means across control and ArchT mice, respectively. n.s. indicates no significance ($p > 0.05$). See Table S8 for the detail of statistical tests. Error bars represent the SEM in all graphs.

**Table S8. Statistical details for Figure S2**

| Fig. | Sample size | Mean and SEM | Statistical test | Values |
|---|---|---|---|---|
| S2A | 6 ArchT mice | No light: 114.29 ± 3.73<br>Yellow: 112.74 ± 5.22 | signed-rank test | W = 10, p = 1.0 |
| S2A | 5 control mice | No light: 109.44 ± 9.83<br>Yellow: 111.74 ± 10.08 | paired t-test | $t_4 = 0.61$, p = 0.57 |
| S2A | 6 ArchT mice<br>5 control mice | ArchT: -0.013 ± 0.032<br>Control: 0.022 ± 0.033 | unpaired t-test | $t_9 = 0.76$, p = 0.47 |
| S2B | 6 ArchT mice | No light: 0.36 ± 0.020 s<br>Yellow: 0.36 ± 0.013 s | signed-rank test | W = 10, p = 1.0 |
| S2B | 5 control mice | No light: 0.40 ± 0.044 s<br>Yellow: 0.40 ± 0.041 s | paired t-test | $t_4 = 0.32$, p = 0.77 |
| S2B | 6 ArchT mice<br>5 control mice | ArchT: 0.0011 ± 0.0200<br>Control: $-5.6 \times 10^{-4}$ ± 0.018 | paired t-test | $t_9 = 0.059$, p = 0.95 |

**Table S9. generalized linear mixed effect model analysis of the number of lever-press in optogenetic inhibition studies**

| Group | Effect | Estimate | 95% CI lower / upper | t-stats | p-values |
|---|---|---|---|---|---|
| ArchT | Intercept | 4.77 | 4.70 / 4.85 | $t_{227} = 124.03$ | $1.5\times10^{-210}$ |
| | Manipulation | -0.0044 | -0.029 / 0.020 | $t_{227} = 0.36$ | 0.72 |
| | Elapsed time | $-6.4\times10^{-5}$ | $-8.2\times10^{-5}$ / $-4.7\times10^{-5}$ | $t_{227} = 7.14$ | $1.2\times10^{-11}$ |
| | Sessions | 0.011 | 0.00021 / 0.022 | $t_{227} = 2.01$ | 0.046 |
| Control | Intercept | 4.74 | 4.58 / 4.90 | $t_{203} = 59.94$ | $4.7\times10^{-131}$ |
| | Manipulation | 0.0172 | -0.00890 / 0.0433 | $t_{203} = 1.30$ | 0.20 |
| | Elapsed time | $-6.7\times10^{-5}$ | $-8.2\times10^{-5}$ / $-5.1\times10^{-5}$ | $t_{203} = 8.45$ | $5.4\times10^{-15}$ |
| | Sessions | 0.017 | 0.0047 / 0.028 | $t_{203} = 2.77$ | 0.0061 |

**Table S10. generalized linear mixed effect model analysis of short IPIs in optogenetic inhibition studies**

| Group | Effect | Estimate | 95% CI lower / upper | t-stats | p-values |
|---|---|---|---|---|---|
| ArchT | Intercept | 0.39 | 0.35 / 0.42 | $t_{227} = 22.099$ | $1.7\times10^{-58}$ |
| | Manipulation | -0.0026 | -0.015 / 0.0097 | $t_{227} = 0.41$ | 0.68 |
| | Elapsed time | $-5.8\times10^{-6}$ | $-1.5\times10^{-5}$ / $3.1\times10^{-6}$ | $t_{227} = 1.3$ | 0.20 |
| | Sessions | -0.0065 | -0.012 / -0.00090 | $t_{227} = 2.3$ | 0.023 |
| Control | Intercept | 0.49 | 0.41 / 0.57 | $t_{203} = 11.66$ | 0.57 |
| | Manipulation | -0.0029 | -0.021 / 0.016 | $t_{203} = 0.31$ | 0.76 |
| | Elapsed time | $-2.1\times10^{-5}$ | $-3.2\times10^{-5}$ / $-1.1\times10^{-5}$ | $t_{203} = 3.92$ | 0.00012 |
| | Sessions | -0.021 | -0.029 / -0.013 | $t_{203} = 4.94$ | $1.6\times10^{-6}$ |
